# Oxygen depletion and nitric oxide stimulate type IV MSHA pilus retraction in *Vibrio cholerae* via activation of the phosphodiesterase CdpA

**DOI:** 10.1101/2021.04.20.440694

**Authors:** Hannah Q. Hughes, Kyle A. Floyd, Sajjad Hossain, Sweta Anantharaman, David T. Kysela, Yuanchen Yu, Michael P. Kappler, Triana N. Dalia, Ram C. Podicheti, Douglas B. Rusch, Yves V. Brun, Stephen C. Jacobson, James B. McKinlay, Fitnat H. Yildiz, Elizabeth M. Boon, Ankur B. Dalia

## Abstract

Bacteria use surface appendages called type IV pili to perform diverse activities including DNA uptake, twitching motility, and attachment to surfaces. Dynamic extension and retraction of pili is often required for these activities, but the stimuli that regulate these dynamics remain poorly characterized. To study this question, we use the bacterial pathogen *Vibrio cholerae*, which uses mannose-sensitive hemagglutinin (MSHA) pili to attach to surfaces in aquatic environments as the first step in biofilm formation. Here, we find that *V. cholerae* cells retract MSHA pili and detach from a surface in microaerobic conditions. This response is dependent on the phosphodiesterase CdpA, which decreases intracellular levels of cyclic-di-GMP (c-di-GMP) under microaerobic conditions to induce MSHA pilus retraction. CdpA contains a putative NO-sensing NosP domain, and we demonstrate that nitric oxide (NO) is necessary and sufficient to stimulate CdpA-dependent detachment. Thus, we hypothesize that microaerobic conditions result in endogenous production of NO (or an NO-like molecule) in *V. cholerae*. Together, these results describe a regulatory pathway that allows *V. cholerae* to rapidly abort biofilm formation. More broadly, these results show how environmental cues can be integrated into the complex regulatory pathways that control pilus dynamic activity and attachment in bacterial species.

## MAIN

Type IV pili are nearly ubiquitous nanomachines in bacteria^1^. These structures promote a wide variety of functions, including the uptake of DNA for horizontal gene transfer^2^, motility on surfaces^3^, and attachment to surfaces^4–6^. Pili are helical filaments composed of a single repeating protein called the major pilin^7^. Major pilin subunits are dynamically assembled into a helical filament on an inner membrane platform protein through the action of a specific motor ATPase protein. This dynamic process of assembly promotes the extension of the pilus filament from the cell surface. Conversely, through the action of an antagonistic ATPase, pilus filaments are depolymerized and major pilin subunits are recycled into the membrane. The dynamic process of pilus disassembly results in the retraction of these structures^8–10^. The dynamic extension and retraction of pili is crucial for their diverse functions. Furthermore, this dynamic activity represents one important mechanism that bacteria use to interact with and respond to their environments. However, the mechanisms by which bacteria regulate pilus dynamic activity in response to environmental cues remains poorly characterized. To address this, we examined pilus dynamic activity of the *V. cholerae* type IV MSHA pilus system. MSHA pili are expressed when *V. cholerae* inhabits its aquatic reservoir and are critical for attachment to abiotic surfaces^11–13^. Here, we uncover an environmental condition that stimulates pilus retraction and characterize the molecular mechanism underlying this response.

## RESULTS

### MSHA pilus retraction is required for cell detachment from well slides

MSHA pili are critical for attachment to abiotic surfaces^12,13^ (**Fig. S1A**), and labelling of MSHA pili^5^ (see Methods) reveals that pilus retraction can promote cell detachment^14^ (**Fig. 1A**). The physiological cues that regulate pilus dynamic activity in most T4P to influence their downstream functions, however, remain unclear. One condition we identified that induces cell detachment in *V. cholerae* is the incubation of cells in the well of a glass slide. We found that cells progressively detach in a wave that propagates from the center of the well to the edge, leaving only cells on the periphery of the well attached to the surface (**Fig. 1B**). To further characterize this response, we first developed an assay that allowed us to simultaneously evaluate the detachment of multiple strains in a consistent environment. This assay, henceforth called a “<u>mix</u>ed <u>d</u>etachment” (MixD) assay, involves mixing equal proportions of two strains that express distinct fluorescent markers, which allows for the subsequent tracking of the detachment of each strain via simple epifluorescence time-lapse microscopy. To validate this approach, we performed a MixD assay of two parent strains expressing either GFP or mCherry (**Fig. 1C**). As expected, the two parent strains detached at similar rates over the course of ~4 minutes. Importantly, this assay allows detachment kinetics of a mutant to be directly compared to the parent strain experiencing the same environmental conditions. Thus, we employed MixD assays to identify factors that regulate this detachment response.

**Fig 1:**
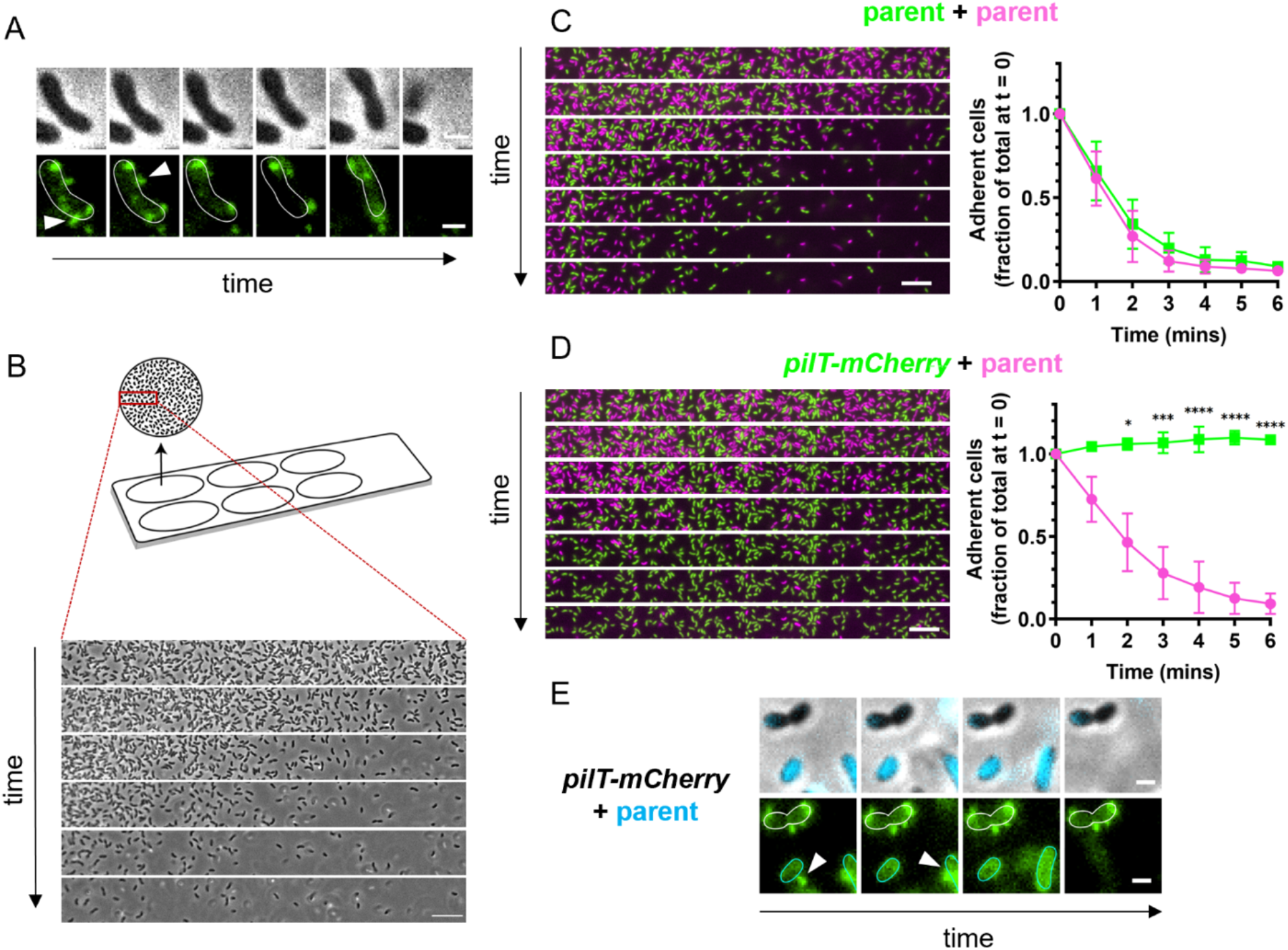
Cells rapidly detach from the surface of a glass well slide in a manner that is dependent on MSHA pilus retraction. (**A**) Representative montage illustrating that MSHA pilus retraction precedes cell detachment. Phase images (top), showing cell boundaries, and FITC channel (bottom) images, showing AF488-mal labelled MSHA pili. White arrows indicate examples where pili retract prior to detachment. 10 second intervals between frames. Scale bar = 1 μm. (**B**) Diagram of experimental setup to observe cell detachment on a glass well slide (top). Representative phase contrast images (bottom) of cell detachment over time. 1-minute interval between frames. Scale bar = 10 μm. (**C-D**) Mixed detachment (MixD) assays of the indicated strains. Cells expressed GFP (green) or mCherry (fuchsia) as indicated by the color-coded genotypes. Representative montages of time-lapse imaging (left) with 1-minute intervals between frames. Scale bar = 10 μm. Quantification of three biological replicates is shown in the line graph (right) and is displayed as the mean ± the standard deviation (SD). Statistical comparisons were made by one-way ANOVA and post-hoc Holm-Šídák test. \**P* <0.05, \*\*\**P* <0.001, \*\*\*\**P* < 0.0001. (**E**) Representative montage of labelled fluorescent MSHA pili in a mixture of a *pilT-mCherry* strain and a CyPet-expressing parent strain. Phase contrast images with overlaid CyPet fluorescence in cyan (top) distinguish the two strains, and FITC fluorescence images in green (bottom) show AF488-mal labelled pili. Parent cells are outlined in cyan and *pilT-mCherry* cells are outlined in white. 20 sec intervals between frames. Scale bar = 1 μm. The PilT-mCherry construct does not exhibit detectable mCherry fluorescence in **D** and **E**.

Because MSHA pilus retraction can promote cell detachment^14^ (**Fig. 1A**), we first wanted to determine if retraction was required for this detachment response. Deletion of the retraction motor gene *pilT* is a commonly employed approach for preventing pilus retraction, which results in hyperpiliation of other T4P^15,16^. For MSHA pili, however, deletion of *pilT* results in a dramatic reduction in pilus biogenesis^14,17^ (**Fig. S2**), for reasons that remain unclear. As an alternative, we took advantage of a natively expressed PilT-mCherry fusion, which we have serendipitously found reduces PilT activity without affecting MSHA pilus biogenesis (**Fig. S2**). We performed a MixD assay of the parent and PilT-mCherry strains and found that while the parent strain detached, the PilT-mCherry strain remained attached throughout the time-lapse (**Fig. 1D**). We further verified that when the parent strain detaches, it is preceded by retraction of the MSHA pilus, and the PilT-mCherry cells that remain attached retain extended MSHA pili (**Fig. 1E**), consistent with a lack of retraction in the latter strain background. This observation indicates that MSHA pilus retraction is required for detachment under these conditions.

### MSHA pilus retraction and cell detachment are dependent on allosteric regulation of MshE activity by c-di-GMP

Transitions between sessile and motile lifestyles are often mediated by the secondary messenger cyclic-di-GMP (c-di-GMP)^18^. Indeed, c-di-GMP regulates *V. cholerae* motility and biofilm formation in part through control of MSHA dynamic activity^14,19^. Specifically, when c-di-GMP levels are elevated through the action of diguanylate cyclases (DGCs), c-di-GMP can bind to the extension motor MshE and allosterically induce pilus extension^20^. Correspondingly, decreased levels of c-di-GMP, which may occur via the activity of phosphodiesterases (PDEs) that cleave c-di-GMP, reduces MshE activity^14,20^ and prevents cell attachment (**Fig. S3**). Thus, we hypothesized that the detachment phenomenon on slides may be caused by a reduction in intracellular c-di-GMP. If so, we would predict that elevating intracellular c-di-GMP should prevent or delay detachment. To test this, we generated strains where we could ectopically express a diguanylate cyclase, *dcpA*^21^, to elevate intracellular c-di-GMP levels (P_BAD_-*dcpA*). Elevated c-di-GMP can also induce expression of the *vps* and *rbm* loci^22^, which encode *Vibrio* polysaccharide and biofilm matrix proteins that can confound attachment in these experiments. Therefore, we deleted both loci (ΔVC0917-VC0939; here called Δ*vps*) in the P_BAD_-*dcpA* background to ensure that attachment in these assays was due to MSHA pili (**Fig. S1B**). When tested in a MixD assay, we found that P_BAD_-*dcpA* Δ*vps* showed significantly delayed detachment compared to the parent when *dcpA* was induced As mentioned above, c-di-GMP is required to allosterically stimulate MshE activity^20^. Previous studies have identified an MshE mutation that maintains activity even in the absence of c-di-GMP (MshE L10A/L54A/L58A, denoted MshE*)^20^. If cell detachment is induced by a reduction in the activity of MshE (due to lowered c-di-GMP levels), we hypothesized that an MshE* mutation should delay or prevent cell detachment. Indeed, in a MixD assay, the MshE* strain showed a significant reduction in detachment compared to the parent (**Fig. 2B**). Consistent with MshE* bypassing c-di-GMP-dependent regulation, attachment of the MshE* strain is not altered even when the PDE *cdgJ* is overexpressed in this background (**Fig S3**). Also, direct labeling of pili in the P_BAD_-*dcpA* Δ*vps* and MshE* strains demonstrated that cells maintained extended MSHA pili, which is consistent with a lack of pilus retraction in these backgrounds (**Fig 2C, D**). Together, these results suggest that cell detachment on slides is caused by MSHA pilus retraction in response to allosteric regulation of MshE by c-di-GMP.

**Fig 2:**
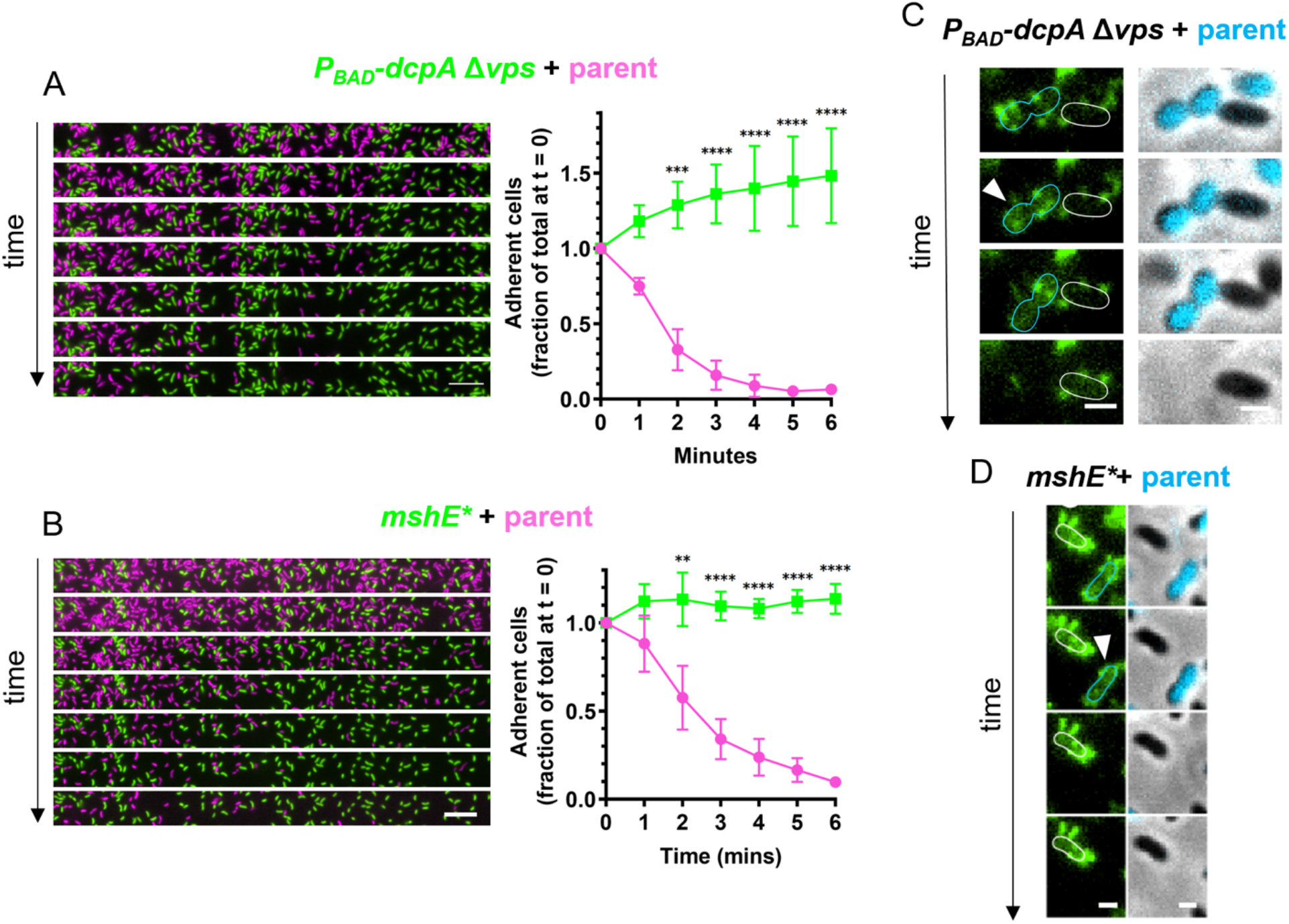
Cell detachment is dependent on allosteric regulation of MshE activity by c-di-GMP. (**A**) MixD assay of *P_BAD_-dcpA* Δ*vps* expressing GFP (green) and a parent strain expressing mCherry (fuschia). *P_BAD_-dcpA* Δ*vps* cells were induced with 0.15% arabinose. Representative montage of time-lapse imaging (left) with 1-minute intervals between frames. Scale bar = 10 μm. Quantification of three biological replicates is shown in the line graph (right) and is displayed as mean + SD. Statistical comparisons were made by one-way ANOVA and post-hoc Holm-Šídák test. \*\**P* <0.01, \*\*\**P* <0.001, \*\*\*\**P* < 0.0001. (**B**) MixD assay as in **A** of *mshE*\* expressing GFP (green) and a parent strain expressing mCherry (fuschia). (**C**) Representative montage of time-lapse imaging of cells with AF488-mal labelled MSHA pili. Cells represent a mixture of *P_BAD_-dcpA* Δ*vps* cells and CyPet-expressing parent cells. *P_BAD_-dcpA* Δ*vps* cells were induced with 0.15% arabinose as in **A**. Phase contrast images with overlaid CyPet fluorescence in cyan (right) distinguish the mixed strains, and FITC fluorescence images in green (left) show labelled pili. Parent cells are outlined in cyan and *P_BAD_-dcpA* Δ*vps* cells are outlined in white. White arrows indicate clear examples where pilus retraction precedes cell detachment. 20 sec intervals between frames. Scale bar = 1 μm. (**D**) Representative montage of time-lapse imaging of cells where pili are fluorescently labelled as in **C**. Cells represent a mixture of *mshE*\* cells and parent cells expressing CFP. Parent cells are outlined in cyan and *mshE*\* cells are outlined in white.

### Cell detachment occurs under microaerobic conditions

Next, we sought to define the environmental stimulus that drives this detachment response. Because well slides were covered with an air-impermeable glass coverslip, the edge of the well is the only place where cells can access O_2_ for cellular respiration. Thus, we hypothesized that O_2_ depletion in the center of the well, which likely occurs as cells respire, is an environmental cue that induces MSHA pilus retraction. To test this, we drilled holes into slides such that the centers of the wells were opened to air. Under these conditions, cells detached as seen previously, except that in the center of the well, cells stayed bound around the hole (**Fig. 3A**). This result is consistent with oxygen availability contributing to persistent cell attachment.

**Fig 3:**
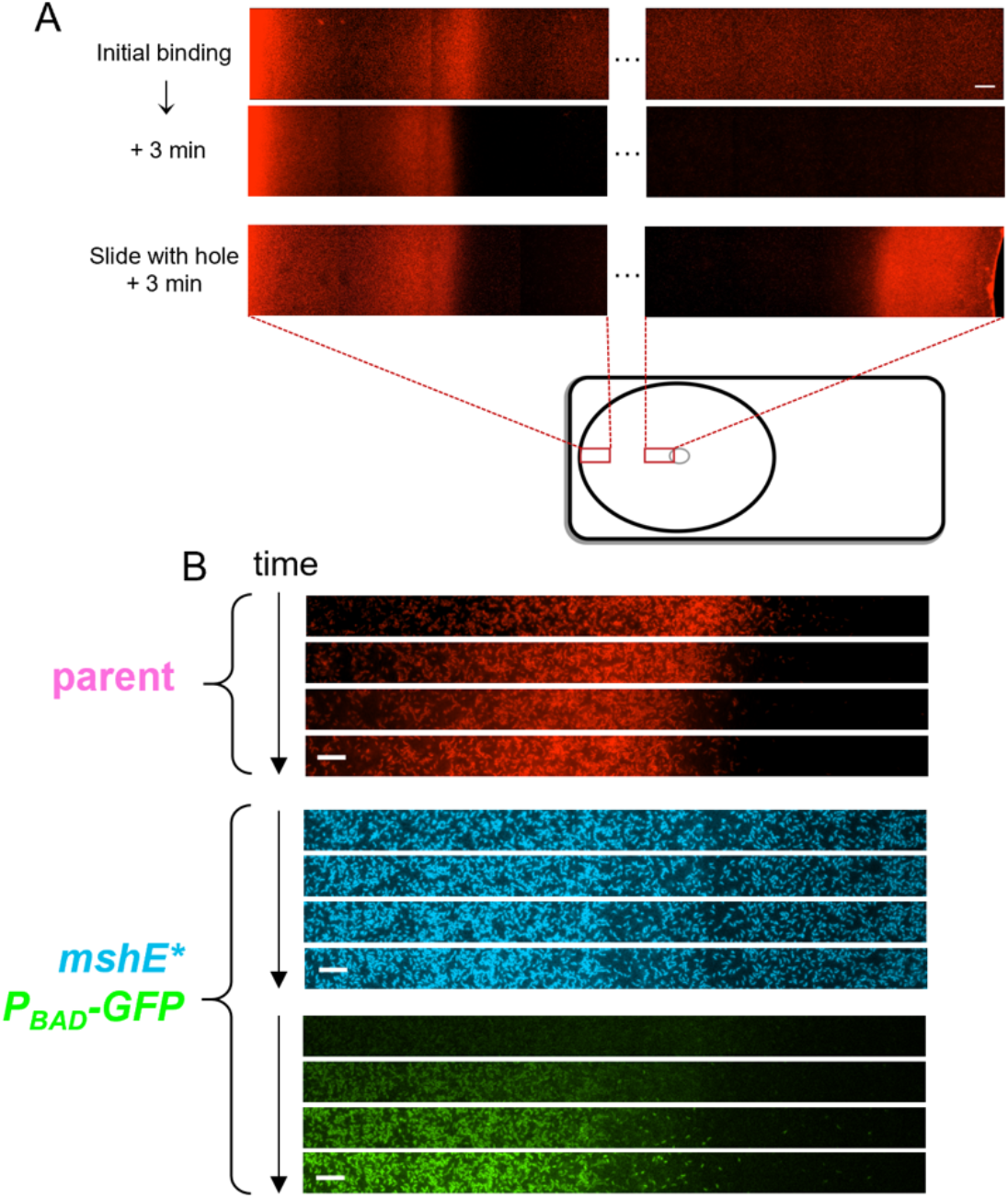
O_2_ depletion induces cell detachment. (**A**) Cell attachment was monitored in well slides by fluorescence microscopy with an mCherry expressing parent strain. Wells either lacked (top and middle) or contained (bottom) a hole in the center of the slide to serve as an air source. Five fields of view are omitted between the edge (left) and center (right) of the well. Scale bar = 50 μm. (**B**) Representative montage of timelapse imaging of a mixed culture of an mCherry expressing parent and a CyPet expressing *mshE*\* strain. The *mshE*\* strain also contains a P_BAD_-*gfp* construct, which serves as a bioreporter for O_2_ in this experiment. Arabinose was added to the mixed culture immediately before imaging. Montages are shown for each fluorescent channel separately with 25-min intervals between frames. Scale bar = 10 μm.

To determine if oxygen depletion correlates with cell detachment, we generated a bioreporter strain that can indirectly indicate oxygen availability. This bioreporter operates on the basis of oxygen being required for the maturation of the GFP chromophore^23^. Specifically, to make an O_2_ reporter we generated a strain in the MshE* background where CyPet (a cyan fluorescent protein derivative) was constitutively expressed and GFP expression was tightly regulated via an arabinose-inducible promoter (P_BAD_-*gfp*). The MshE* mutation prevents detachment in this experimental setup (as in **Fig. 2C**), the constitutive expression of CyPet allows for easy cell tracking, and the addition of arabinose allows for the *de novo* production of GFP. To track the kinetics and boundaries of cell detachment, we mixed this O_2_ reporter strain with a parent strain constitutively expressing mCherry. In this experiment, arabinose was added immediately before strains were placed on well slides, thereby inducing expression of GFP protein in all CyPet labeled cells. However, GFP would only mature into its fluorescent form where sufficient oxygen was present. This process was monitored by epifluorescence time-lapse microscopy. As expected, the parent strain rapidly detached while the MshE* strain remained attached throughout the time-lapse. Notably, only cells at the edge of the well exhibited GFP fluorescence (**Fig. 3B**), which is consistent with a lack of GFP maturation in the center of the well due to oxygen depletion. Interestingly, the boundary of GFP fluorescence was similar to the boundary of cell attachment observed in the parent strain (**Fig. 3B**). Although not necessarily expected, this result indicated that the threshold of oxygen required for cell attachment is similar to the threshold required for GFP maturation. Together, these results strongly suggest that O_2_ depletion is an environmental cue that induces MSHA pilus retraction and subsequent cell detachment.

Thus far, our data suggest that O_2_ depletion induces MSHA pilus retraction in a manner that depends on a decrease in cellular c-di-GMP. We, therefore, hypothesized that *V. cholerae* carries a PDE that, during O_2_ depletion, decreases c-di-GMP levels, triggering MSHA pilus retraction and cell detachment. We would expect a mutant lacking this putative PDE to exhibit enhanced adherence under microaerobic conditions (akin to the MshE* mutant). To test this hypothesis, we first established conditions that would allow us to study the effect of oxygen depletion in a larger culture format. To accomplish this, we allowed cells to bind to the sides of either open, rolling glass tubes (aerobic; “+air”), or sealed, static argon-flushed glass tubes (microaerobic; “-air”) (see Methods). We confirmed that these culture conditions recapitulated the binding phenotypes seen in the MixD assays, where the parent strain bound poorly under microaerobic conditions and the MshE* mutant bound equally well under both aerobic and microaerobic conditions (**Fig. S4A**). We also directly measured oxygen levels to confirm that our culture conditions generated aerobic and microaerobic environments, as expected (**Fig. S4B, C**).

### Cell detachment requires the phosphodiesterase CdpA

We then carried out a genetic selection to isolate transposon mutants that exhibited enhanced adherence under microaerobic conditions, thereby identifying factors, like the hypothesized PDE, that might be required to stimulate cell detachment. Following the genetic selection, we isolated 10 colonies and sequenced the Tn-genomic junction in each. All mutants contained Tn insertions (9 distinct mutations) in *cdpA* (VC0130) (**Fig. S5A**). CdpA is a functional PDE that contains a NosP sensory domain^24,25^. In a MixD assay, the Δ*cdpA* mutant exhibited significantly delayed detachment compared to the parental strain (**Fig. S5B**). Importantly, this phenotype could be complemented *in trans* with a single copy of *cdpA* integrated at an ectopic location under the control of its native promoter (Δ*cdpA* Δ*lacZ*::*cdpA^WT^*) (**Fig. 4A**). These results confirm that CdpA plays an important role in cell detachment under microaerobic conditions.

**Fig 4:**
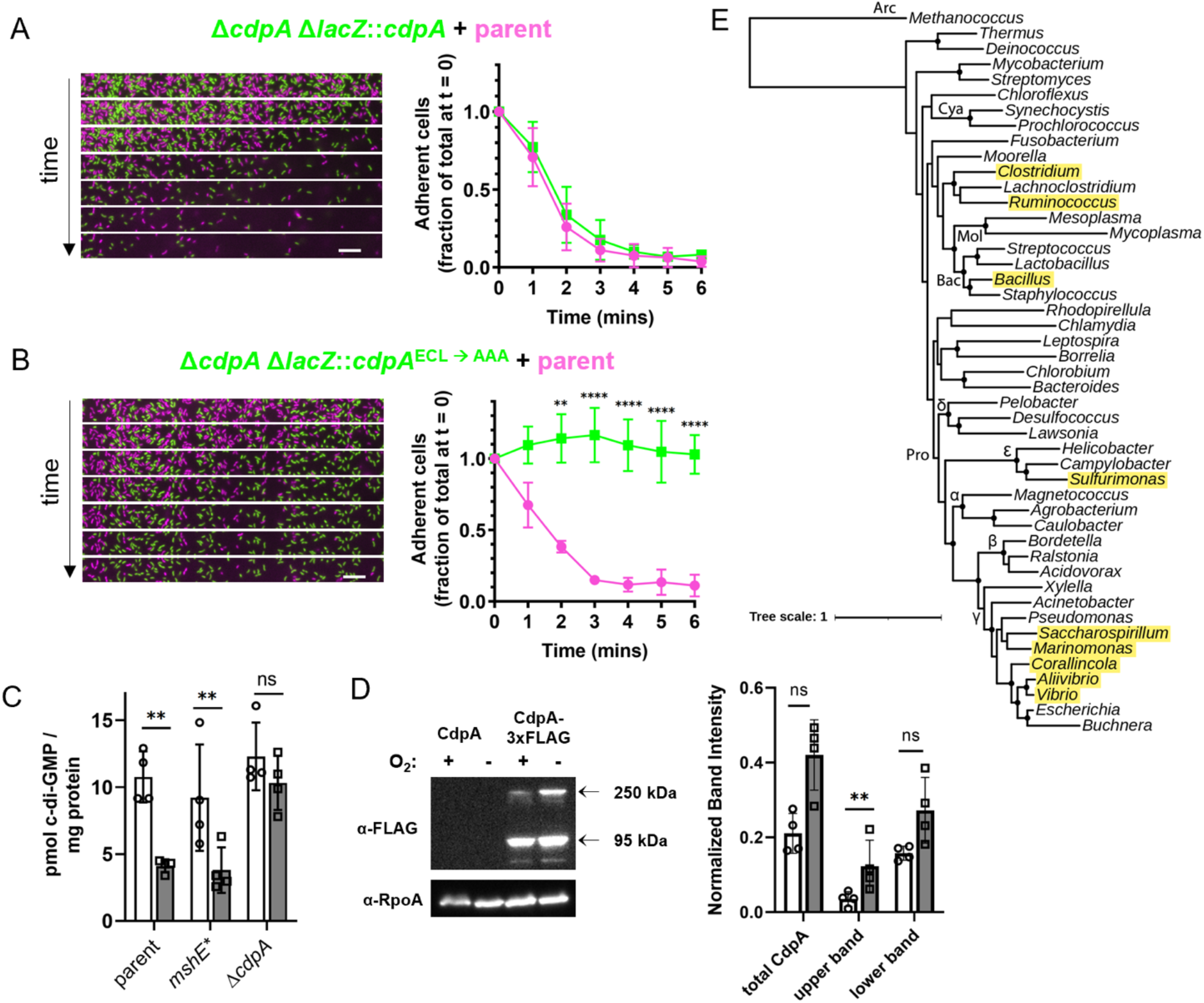
CdpA promotes cell detachment in microaerobic conditions via its PDE activity. (**A**-**B**) MixD assays of the indicated strains. Cells expressed GFP (green) or mCherry (fuchsia) as indicated by the color-coded genotypes. Complemented strains are denoted by “Δ*lacZ*::*cdpA*”, indicating that a copy of *cdpA* along with its native promoter were introduced at an ectopic site on the chromosome. Representative montage of time-lapse imaging (left) with 1-minute intervals between frames. Scale bar = 10 μm. Quantification of three biological replicates is shown in each line graph (right). (**C**) Quantification of intracellular c-di-GMP concentrations in the indicated strains. Data are from four independent biological replicates. Open bars represent samples from aerobic cells while shaded bars indicate samples from microaerobic cells. (**D**) Representative western blot (right) and quantification (left) measure CdpA protein levels under aerobic and microaerobic conditions. Open bars represent aerobic cells and shaded bars represent microaerobic cells. Band intensities were normalized to the RpoA loading control (visualized on a separate blot). Data are from four independent experiments. Statistical comparisons were made on log-transformed data. (**E**) Conservation of CdpA across bacterial species. Estimated maximum likelihood phylogeny of the indicated microbes based on a concatenated alignment of 36 conserved proteins identified from whole genome sequences. Genera with members that contain a CdpA homolog are highlighted in yellow. For a complete list of species that contain a CdpA homolog see **Table S1**. Major taxa are labeled along their nodes. Pro: Proteobacteria (Greek letters indicate subdivisions); Bac: Bacilli; Mol: Mollicutes; Cya: Cyanobacteria; Arc: Archaea. Scale bar indicates distance. All graphs display mean ± SD. Statistical comparisons were made by one-way ANOVA and post-hoc Holm-Šídák test. \**P* <0.05, \*\**P* <0.01, \*\*\*\**P* < 0.0001.

Because CdpA is a PDE, we hypothesized that it reduced c-di-GMP under microaerobic conditions to promote MSHA pilus retraction and subsequent cell detachment. To test this, we first mutated the EAL domain required for PDE activity in *cdpA* (*cdpA* has ECL instead of the canonical EAL within this domain). In MixD assays, the Δ*cdpA* Δ*lacZ*::*cdpA*^ECLàAAA^ strain phenocopied the Δ*cdpA* mutant (**Fig. 4B**), indicating that the PDE activity of CdpA plays a critical role in promoting cell detachment. Consistent with this result, we found that intracellular levels of c-di-GMP (measured by LC-MS/MS) were reduced under the microaerobic conditions that stimulate cell detachment in a CdpA-dependent manner (**Fig. 4C**, **S4A**).

Based on the data described above, CdpA likely degrades c-di-GMP specifically under microaerobic conditions. What remains unclear is the mechanism by which CdpA activity is stimulated. One possibility is that upon a decrease in oxygen, CdpA protein levels are increased. To test this hypothesis, we generated a functional *cdpA*-3xFLAG strain (**Fig. S5C**) and assessed protein levels by western blot. This revealed that there was a slight, albeit not significant, increase in total CdpA levels when cells were incubated microaerobically (**Fig. 4D**). We did find that the intensity of a high molecular weight CdpA band is increased in western blots when cells are incubated microaerobically (**Fig. 4D**). This band may represent an active CdpA complex. Regardless, because total CdpA levels are not dramatically altered in these experiments, we hypothesized that CdpA activity is likely modulated via a post-translational mechanism.

### NO is necessary and sufficient to promote MSHA pilus retraction and cell detachment in a CdpA-dependent manner

Other oxygen-responsive PDEs and DGCs have been characterized, often sensing oxygen through a heme-binding PAS or globin domain^26–29^. CdpA lacks these domains, and instead encodes a NosP domain, which has recently been shown to bind heme in a manner that alters its PDE activity^25^. NosP domains have previously been implicated in sensing NO^30^, and bacterial NO production generally occurs under microaerobic conditions^31,32^. Thus, we hypothesized that microaerobic conditions may indirectly induce MSHA retraction by promoting NO production, which induces CdpA activity. Therefore, we tested whether NO was necessary and sufficient to induce CdpA-dependent detachment. To determine if NO was necessary for detachment, we incubated cells on sides with the NO scavenger 2-phenyl-4,4,5,5,-tetramethylimidazoline-1-oxyl 3-oxide (PTIO). In the presence of PTIO, parent cells remained attached to the surface in a manner that resembled the Δ*cdpA* mutant (**Fig. 5A**). To exclude the possibility that PTIO was non-specifically scavenging reactive oxygen species (ROS)^33^, we also assessed the impact of the ROS-specific scavenger Tiron. Tiron did not inhibit detachment of the parent strain, suggesting that PTIO inhibits cell detachment by specifically scavenging NO, or a related molecule (**Fig. 5A**). To test if NO was sufficient to induce detachment, we incubated cells with the NO donor diethylamine-NONOate (DEA-NONOate) and assessed cell attachment under aerobic conditions. We found that parent cells attached poorly in the presence of DEA-NONOate, whereas attachment of the Δ*cdpA* mutant was unaffected (**Fig. 5B**). Furthermore, incubation with DEA-NONOate resulted in MSHA pilus retraction in a CdpA-dependent manner (**Fig. 5C**). Together, these observations are consistent with NO being necessary and sufficient to induce MSHA pilus retraction via stimulating CdpA activity.

**Fig 5:**
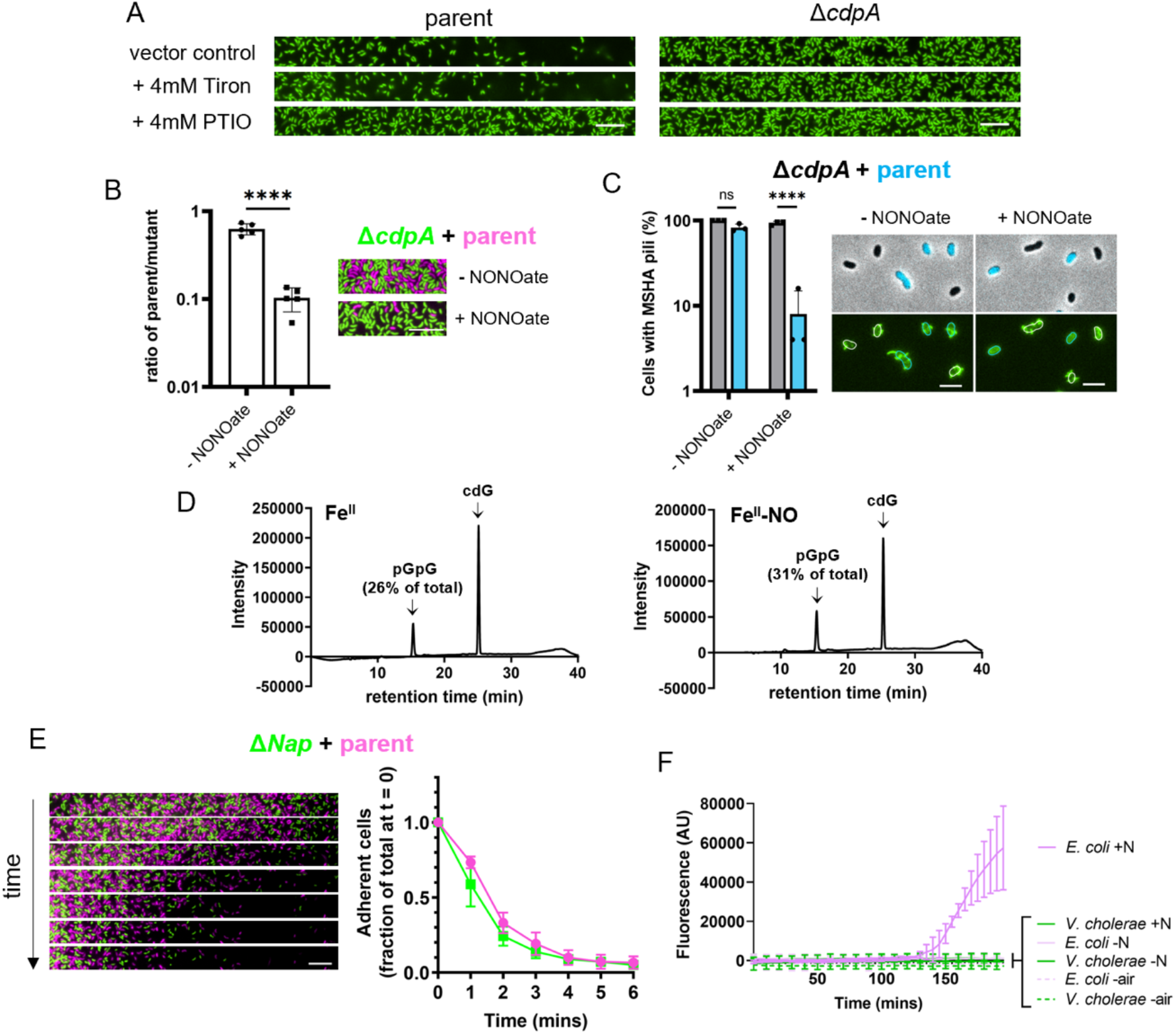
Nitric oxide stimulates detachment in a CdpA-dependent manner. (**A**) Representative detachment images of GFP-expressing parent or Δ*cdpA* strains after incubation with Tiron (4 mM), PTIO (4 mM), or DMSO as a vector control. Scale bar = 10 μm. (**B**) Quantification (left) of initial attachment for a 1:1 mixture of the GFP-expressing parent (green) and mCherry-expressing Δ*cdpA* mutant (fuchsia) when incubated in the presence or absence of 200 μM DEA-NONOate, as indicated. Ratio is determined by dividing the number of attached parent cells (green) by the number of attached Δ*cdpA* mutant cells (fuchsia) as depicted in the representative image on the right. Data are from five independent experiments with at least 270 cells analyzed per replicate. Statistical comparison made by t-test. Scale bar = 10 μm. (**C**) Quantification (left) of parent (blue) or Δ*cdpA* (gray) piliation after incubation in the presence or absence of 200 μM DEA-NONOate. Fraction is determined by dividing number of cells with pili by the total number of cells counted for that strain for that condition. Data are from three independent experiments with at least 50 cells analyzed per replicate, per condition. Representative images (right) of cells with AF488-mal labelled MSHA pili. Cells represent a mixture of Δ*cdpA* cells and CyPet-expressing parent cells. Phase contrast images with overlaid CyPet fluorescence in cyan (top) distinguish the mixed strains, and FITC fluorescence images in green (bottom) show labelled pili. Parent cells are outlined in cyan and Δ*cdpA* cells are outlined in white. Scale bar = 3 μm. (**D**) Representative HPLC analysis of purified CdpA PDE activity, indicating the amount of c-di-GMP converted into pGpG. Reactions containing c-di-GMP and purified CdpA with Fe^II^ heme (left) or purified CdpA with Fe^II^-NO (right) were incubated for 24 hr at room temperature. (**E**) MixD assays of parent and Δ*Nap* strains. Cells expressed GFP (green) or mCherry (fuchsia) as indicated by the color-coded genotypes. Representative montage of time-lapse imaging (left) with 1-minute intervals between frames. Scale bar = 10 μm. Quantification of three biological replicates is shown in the line graph (right). (**F**) DAR 4M-AM measurement of NO production in cells. Mean ± SD of fluorescence values for three independent, biological replicates are graphed. Dashed lines indicate samples incubated microaerobically. +/-N indicates incubation with or without 20 mM nitrate. The “-air” condition denotes the microaerobic conditions that stimulate *V. cholerae* detachment. Data in panels **B** and **D**-**F** display the mean ± SD. Statistical comparisons were made by one-way ANOVA and post-hoc Holm-Šídák test. ns = not significant, \*\*\*\**P* < 0.0001.

Next, we wanted to determine how widespread this CdpA-dependent activity may be. Phylogenetic analysis revealed that CdpA is largely conserved among the *Vibrionaceae* and is also prevalent in *Aliivibrio* species (**Fig. S6, S7**). Based on sequence similarity and presence within the *Vibrio* and *Allivibrio*, we conclude that these are homologs with a conserved functional role. Furthermore, we found *cdpA* homologs in phylogenetically distant species (e.g., *Clostridium* and *Bacillus*) (**Fig. 4E, S8, Table S1**). Thus, it is tempting to speculate that CdpA may be a widely conserved, horizontally transferred NO-responsive PDE.

Above, our data suggest that CdpA may be post-translationally regulated. Because the heme-bound NosP domain of CdpA is implicated in directly reacting with NO, we also assessed the impact of NO on the PDE activity of purified heme-bound CdpA. This assay showed that there was a slight, but inconsistent increase in PDE activity for Fe^II^-NO CdpA compared to CdpA bound to heme alone (**Fig. 5D**). This discrepancy in NO-mediated activity compared to the *in vivo* data may be due to limitations of the *in vitro* setup. While the heme moiety used for this experiment (heme b) matches what was observed natively bound to other NosP-containing proteins purified from *E. coli*^30^, CdpA may bind a different heme moiety *in vivo*. Alternatively, there may be an unidentified cofactor required for CdpA activity which is present *in vivo* but lacking in our *in vitro* assays. In support of this hypothesis, we show that CdpA PDE activity is very poor *in vitro* (requires overnight incubations) (**Fig. 5D**), despite the fact that it exerts its effect *in vivo* within a matter of minutes (**Fig. 4**).

A major tenet of our model is that *V. cholerae* endogenously generates NO under microaerobic conditions to stimulate CdpA activity. An important caveat to this conclusion, however, is that *V. cholerae* lacks the canonical denitrification or dissimilatory nitrate reduction to ammonia (DNRA) pathways that allow many bacterial species to generate NO. *V. cholerae* encodes a nitrate reductase (in the *Nap* operon) that converts nitrate to nitrite, but it lacks homologs of other denitrification or DNRA enzymes^34,35^. Although *V. cholerae* encounters NO during infection^36–38^, it is not thought to generate NO endogenously. MixD analysis of a Δ*Nap* mutant^39^ (**Fig. S9A**) was indistinguishable from the parent (**Fig. 5E**), suggesting that nitrate reductase activity is not required to generate a molecular cue for cell detachment. Thus, if NO is being generated, it must be through a pathway previously uncharacterized in *V. cholerae*.

To test whether *V. cholerae* was actually generating NO, we incubated cells with diaminorhodamine-4M acetoxymethyl ester (DAR-4M AM), an NO-specific fluorescent probe. Using DAR-4M AM, we readily measured NO produced by DEA-NONOate (**Fig. S9B**) and *E. coli*, which has an intact DNRA pathway, but were unable to detect NO production from *V. cholerae*, even when cells were incubated under the microaerobic conditions that induce cell detachment (**Fig 5F**). Importantly, NO was still detected by DAR-4M AM when cells were mixed with DEA-NONOate, albeit to a reduced level. These data suggest that cells may slightly decrease the threshold for NO detection by DAR-4M AM by acting as an NO “sponge”. Thus, it remains feasible that concentrations of NO lower than our limit of detection in the presence of cells (i.e., 40 μM DEA-NONOate) are generated in *V. cholerae*. Furthermore, there is evidence that extracellular NO may not equilibrate with cytoplasmic levels^40^, allowing for the possibility that NO generated endogenously by *V. cholerae* may be at levels too low to be measured by this technique. Alternatively, *V. cholerae* may be generating a distinct, NO-like molecule that promotes cell detachment under microaerobic conditions. This could explain why NO was not able to strongly stimulate CdpA PDE activity *in vitro* (**Fig. 5D**) and why NO was not detected by DAR-4M AM in cells incubated microaerobically (**Fig. 5F**). Defining the precise environmental signals and cofactors required to stimulate CdpA PDE activity will be a focus of future work. While our data indicate that NO is sufficient to stimulate cell detachment in *V. cholerae* (**Fig. 5B**), the means by which NO (or an NO-like molecule) is endogenously generated remains unclear.

## DISCUSSION

While the frequency and timing of pilus extension and retraction are crucial to their function, many questions remain about how pilus dynamics are regulated. Some progress has been made on this front, including characterization of protein regulators that impact extension^41–43^ and the impact of spatial organization of the extension and retraction motors^44–46^. Our study highlights another mode of regulation for pilus dynamic activity, wherein environmental regulatory cues rapidly induce pilus retraction through modulation of a secondary messenger (c-di-GMP) that is required for pilus biogenesis (**Fig. 6**).

**Fig 6:**
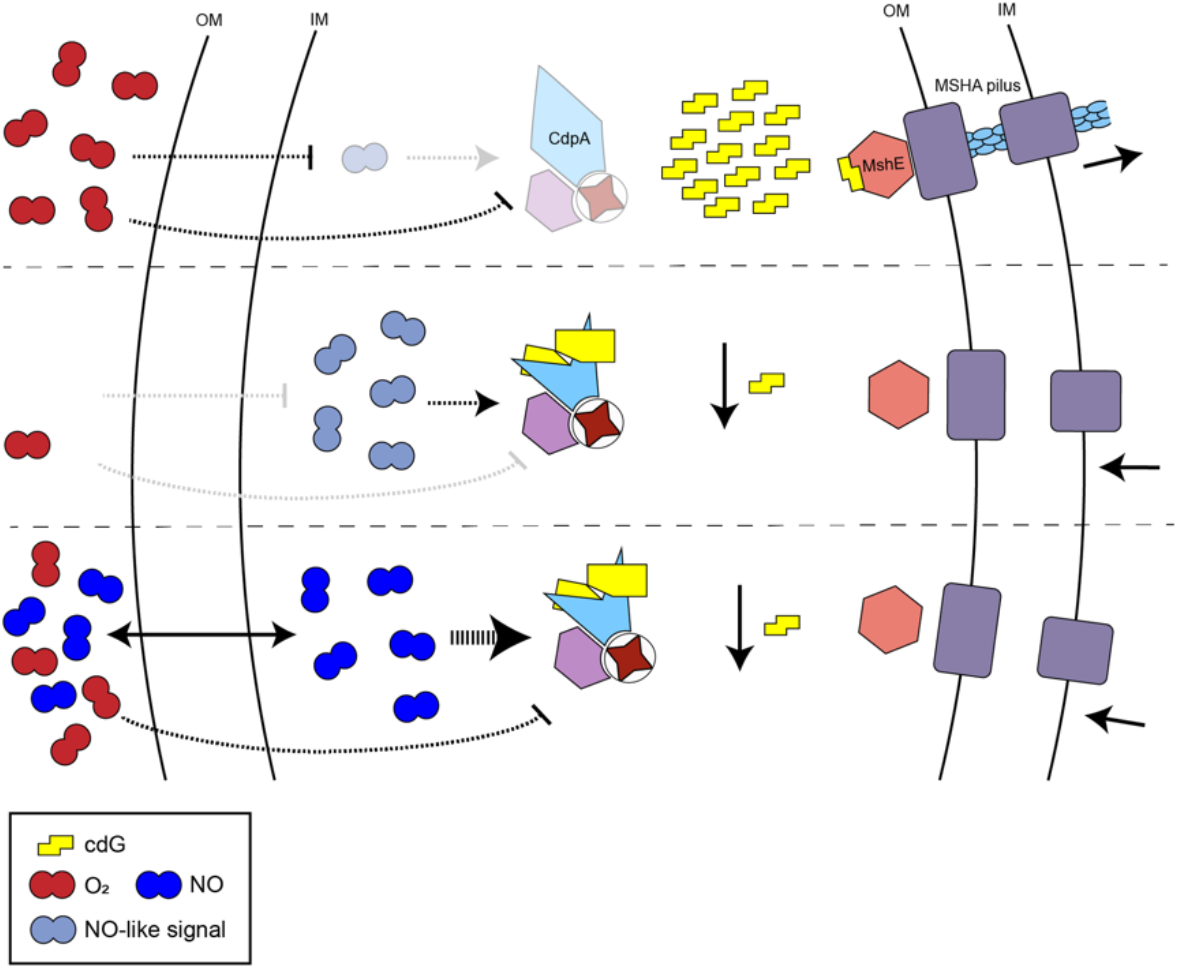
Proposed model for CdpA-dependent induction of MSHA pilus retraction. Under aerobic conditions (top), high oxygen levels inhibit the production of an endogenous NO-like molecule. As a result, CdpA remains inactive, and intracellular c-di-GMP levels stay elevated. This stimulates MshE activity, which causes extension of the MSHA pili and promotes attachment of cells to a surface. Under microaerobic conditions (middle), cells endogenously generate low quantities of NO (or an NO-like signal) through an unknown mechanism. Low oxygen and the NO-like signal, along with a putative cofactor (depicted in lavender), stimulate the PDE activity of CdpA. This decreases the intracellular concentration of c-di-GMP, prompting the retraction of MSHA pili and detachment of cells from the surface. Exogenously added NO (bottom) is sufficient to stimulate CdpA activity even under aerobic conditions (possibly because stimulation by NO is stronger than inhibition by O_2_), which results in decreased c-di-GMP, retraction of MSHA pili, and cell detachment.

CdpA is only one of at least 62 enzymes that contribute to c-di-GMP production and/or degradation in *V. cholerae*^47^. These diverse DGCs and PDEs are believed to respond to distinct signals/cues, thus allowing cells to modulate c-di-GMP under different environmental conditions^18^. While inducing signals have been determined for some of these enzymes^48–50^, the complex regulatory interplay between them is largely uncharacterized and many of the environmental cues required to stimulate their activities remain unclear. Here, we uncover that CdpA responds to NO (and/or an NO-like signal) that is endogenously generated under microaerobic conditions to reduce intracellular c-di-GMP. One outcome for this response is the retraction of MSHA pili, which results in cell detachment.

MSHA pili are principally required by *V. cholerae* to form biofilms in the aquatic environment. Retraction of MSHA pili under microaerobic conditions and/or in the presence of NO, however, may also be biologically relevant when this facultative pathogen infects its human host. Expression of MSHA pili during infection is known to attenuate *V. cholerae* due to the propensity of these appendages to bind secretory IgA in the gut, which promotes their clearance from the small intestine^51^. Indeed, the expression of the MSHA locus is downregulated during infection^51,52^. As the small intestine is a low-oxygen, high-NO environment^53,54^, we speculate that these cues could represent an important trigger to induce rapid MSHA pilus retraction, contributing to *V. cholerae’s* ability to evade the host innate immune response during infection.

## ACKNOWLEDGEMENTS

We would like to thank Clay Fuqua, Julia van Kessel, Tuli Mukhophadyay, and Ilana Heckler for helpful discussions. This work was supported in part by grants R35GM128674 (to ABD), R01AI102584 (to FHY), GM118894 (to EMB), and R01GM113121 (to SCJ) from the National Institutes of Health. YVB is supported by a Canada 150 Research Chair from the Canadian Institutes of Health Research.

## METHODS

### Bacterial strains and culture conditions

*Vibrio cholerae* E7946 was used throughout the study. Cultures were routinely grown in LB broth (Miller) and plated on LB agar at 30°C. Descriptions of all strains used in this study are found in **Supplemental Table 2**. Where appropriate, media were supplemented with kanamycin 50 μg/mL, trimethoprim 10 μg/mL, spectinomycin 200 μg/mL, chloramphenicol 1 μg/mL, carbenicillin 100 μg/mL, or zeocin 100 μg/mL.

### Construction of mutant strains

Mutant constructs were generated by splicing-by-overlap extension PCR as described previously^55^. Natural transformation and cotransformation were used to introduce mutant constructs into chitin-induced competent *V. cholerae* as described previously^56,57^, and mutations were confirmed by PCR and/or sequencing. All ectopic expression constructs were derived from previously published strains^58^. The specific primers used to generate mutant constructs can be found in **Supplemental Table 3**.

### Transposon mutagenesis and selection

Transposon mutant libraries were generated with the mini-Tn10 delivery plasmid pDL1098 as previously described^59^. To carry out the genetic selection, the transposon mutant library was grown to OD_600_ of ~2.5 at 30°C shaking. Cultures were transferred to a stoppered argon-flushed glass tube and incubated statically at room temperature for 3 hours. The tube was then rinsed 5 times with LB to remove unbound cells. Bound cells were removed from the walls of the glass tube by vortexing vigorously with fresh LB. Samples were taken before and after incubation in the argon tube for quantitative plating. Cells retrieved after binding were used to inoculate the next round of selection. Six rounds of selection were carried out, at which point the binding frequency (CFU after incubation divided by CFU before incubation) matched the level of an MshE* control. Then, a sample from the endpoint population was isolation streak plated and ten individual colonies were picked. The transposon-genomic junction in these colonies was amplified via inverse PCR and sent for Sanger sequencing to identify the gene mutated.

### Microscopy and image analysis

All phase contrast and fluorescence images were obtained with a Nikon Ti-2 microscope with a Plan Apo ×60 objective, a green fluorescent protein filter cube, a yellow fluorescent protein filter cube, a cyan fluorescent protein filter cube, an mCherry filter cube, a Hamamatsu ORCAFlash 4.0 camera, and Nikon NIS Elements imaging software. In strains expressing both GFP and CyPet, the GFP signal was imaged with the yellow fluorescent protein filter cube to minimize spectral overlap. Images were prepared with Fiji^60,61^. For experiments where bound cells needed to be quantified, the distinct fluorescence of each cell in the mixture was used to quantify the adherence of each cell type within each frame of the time lapse. MicrobeJ^62^ was used to automate this analysis on regions that contained ~300 total cells.

### MixD analysis

Cells were first grown in LB + 100 μM IPTG (+ 0.15% arabinose, where applicable) to OD_600_ of ~2.5 at 30°C rolling. Cultures were mixed at a 1:1 ratio, loaded onto a multiwell glass slide, and then covered with a glass coverslip. Each culture in these assays expressed a distinct fluorescent protein to help track each individual cell type by microscopy. For each sample, we captured a time-lapse series where phase contrast and fluorescence images were captured once every minute. Replicates were normalized by aligning the first time point to the frame immediately prior to the first recorded decrease in bound cells in the parent strain (i.e., when detachment occurs in the parent strain). Attached cell counts in subsequent frames were divided by the number of cells bound in the first frame.

### Assessing cell attachment

To assess initial adherence of bacteria, cells were grown in LB (+ 100 μM IPTG and/or 0.15% arabinose, where applicable) to OD_600_ of ~2.5 at 30°C rolling. To assess the adherence of a single strain, a pure culture was loaded onto a multiwell glass slide and imaged by phase contrast microscopy. For experiments comparing initial attachment of P_BAD_-*cdgJ* with parent, cells were mixed and added to a coverslip. Cells were allowed to bind aerobically prior to rinsing with instant ocean medium (7 g/L; Aquarium Systems). Rinsed spots were covered with a 0.2% gelzan pad and imaged. Binding frequency was determined by dividing cell counts for each mutant by the cell count of the parent strain.

To test the effect of access to air on cell adherence, ~2 mm holes were sandblasted in the center of wells of a multiwell glass slide. Cells were grown overnight in LB + 100 μM IPTG at 30°C rolling. Cells were loaded onto a multiwell glass slide (either with or without a hole), covered with a glass coverslip, and imaged.

For large-scale binding assays, cells were grown to OD_600_ of ~2.5 at 30°C shaking and loaded into either an open glass tube (aerobic) or into a stoppered argon-flushed glass tube (microaerobic). Aerobic tubes were incubated on a roller drum, while microaerobic tubes were incubated statically. All cultures were incubated at room temperature for 3 hours and cells were allowed to bind to the tubes. After incubation, tubes were dumped and rinsed 5 times with fresh LB to remove unbound cells. Bound cells were then removed from the walls of the glass culture tube by vortexing for 1 min with 3 mL fresh LB. The cell slurry was then centrifuged and concentrated prior to quantitative plating. The binding frequency was calculated for each sample by dividing the CFU after incubation (bound cells) by the CFU before incubation (total cells). Five biological replicates were carried out for each strain.

For assessment of binding during PTIO scavenging, cells were first grown in LB + 100 μM IPTG to an OD_600_ of ~2.5 at 30°C shaking. PTIO or Tiron (or an equivalent volume of DMSO vector control) were added to aliquots of these cultures to a final concentration of 4 mM. Cells were loaded onto a multiwell slide, covered with a coverslip, and incubated for 5 minutes prior to imaging.

For assessment of binding during incubation with DEA-NONOate, cells were first grown in LB + 100 μM IPTG to an OD_600_ of ~2.5 at 30°C shaking. Cells were then mixed with DEA-NONOate (or an equivalent volume of 0.1N NaOH vector control) to a final concentration of 200 μM. Culture was added to a coverslip, and cells were allowed to bind aerobically prior to rinsing with instant ocean medium. Rinsed spots were covered with a 0.2% gelzan pad and imaged. Binding frequency was determined by dividing cell counts of the parent strain by the cell count of the mutant strain.

### Visualization of labelled pili

To visualize pili in the context of a MixD assay, cells were first grown in LB + 100μM IPTG (+0.15% arabinose, where applicable) to OD_600_ of ~2.5 at 30°C rolling. Then, ~10^9^ cells were spun down at 18,000 x g for 1 min and resuspended in instant ocean medium. Cells were then incubated with 25 μg/mL AF488-mal for 10 min at room temperature. Cells were washed twice to remove excess dye and resuspended in instant ocean medium. After labelling, the strain of interest and the parent strain were mixed at a ratio of 1:4, loaded onto a multiwell glass slide, covered with a glass coverslip, and imaged. For each sample, we captured a time-lapse series where phase contrast and fluorescence images were captured every 5 seconds.

To visualize pili in cultures incubated with DEA-NONOate (**Fig. 5C**), cells were labelled with AF488-mal as above, except cells were resuspended in spent LB medium. A 1:1 mixture of labelled cells was incubated with DEA-NONOate (or an equivalent volume of 0.1N NaOH vector control) to a final concentration of 200 μM. Samples were allowed to incubate at room temperature for 5 minutes and were then fixed by addition of paraformaldehyde to a final concentration of 2.5% and incubation at room temperature for 15 mins. Cells were then spun down at 18,000 x g for 1 min and resuspended in instant ocean medium. Samples were loaded onto a coverslip, covered with a 0.2% gelzan pad, and imaged. Twenty-five cells of each strain were randomly chosen and scored for the presence or absence of pili for each condition within each replicate.

### Oxygen quantification

Oxygen levels in glass tubes, with and without culture, were measured with the non-invasive fiber optic PreSens Fibox 4 oxygen meter. Three biological replicates were measured for each scenario. A time course of culture-dependent oxygen-depletion was obtained by taking a measurement once per second after tubes were removed from the roller drum.

To indirectly detect the presence of oxygen with GFP maturation as a bioreporter, cells were first grown in LB + 100 μM IPTG to OD_600_ of ~2.5 at 30°C rolling. Parent (expressing mCherry) and P_BAD_-GFP strains (expressing CyPet) were mixed in a 1:1 ratio and then supplemented with 0.2% arabinose. Immediately thereafter, 2.5 μL of the mixture was added to a multiwell slide, covered with a coverslip, and imaged. Time-lapse series were recorded, capturing phase contrast and fluorescence images once every 5 minutes.

### c-di-GMP quantification

Cells were grown aerobically and microaerobically exactly as described above for large-scale binding assays. Then, cultures were put on ice for 20 minutes. Equivalent volumes of cultures (6.4 mL of culture at OD_600_ ~2.5) were spun down for 15 min at 8,000 x g, at 4° C. Cell pellets were frozen at −80° C until all samples were prepared for c-di-GMP quantification.

Extraction of intracellular c-di-GMP was performed as previously described^63^. Cell pellets were re-suspended in 1 mL of c-di-GMP extraction buffer (40% acetonitrile, 40% methanol, 0.1% formic acid, 19.9% water), vortexed for ~30 seconds, and then incubated on ice for 15 min. Extracted samples were then centrifuged at 16,000 x *g* for 5 min at 4° C, and 800 μl of the c-di-GMP-containing supernatant was removed to a new 1.5mL tube and dried under vacuum. Finally, the dried samples were re-suspended in 50 μl of 184 mM NaCl and analyzed by liquid chromatography-tandem mass spectrometry (LC-MS/MS).

LC-MS/MS analysis was performed in conjunction with the UCSC Chemistry and Biochemistry Mass Spectrometry facility, using a Thermo LTQ-Orbitrap Velos Pro mass spectrometer coupled to a UPLC (Thermo). For UPLC separation, a Synergin Hydro 4u Fusion-RP 80A column (150 mm x 2.00 mm diameter; 4-μm particle size) (Phenomenex, Torrance, CA) was used. Solvent A was 0.1% acetic acid in 10 mM ammonium acetate, solvent B was 0.1% formic acid in methanol. The gradient used was as follows: time (t) = 0–4 min, 98% solvent A, 2% solvent B; t = 10–15 minutes, 5% solvent A, 95% solvent B.

The c-di-GMP levels were calculated from a standard curve generated from pure c-di-GMP suspended in 184 mM NaCl (Sigma) at 0, 25, 50, 100, 500, 2000 nM. The assay is linear from 0 to 2000 nM with an R^2^ of 0.999. Finally, c-di-GMP levels were normalized to total protein per mL of culture. Cell pellets from 1 mL of each sample were lysed with 2% sodium dodecyl sulfate (SDS), and total protein levels of each sample were determined by BCA assay (Thermo). LC-MS/MS analysis was carried out on four biological replicates of each strain to quantify the abundance of c-di-GMP in each sample.

### Western blots

Cells were grown aerobically and microaerobically exactly as described above for large-scale binding assays. Cells were spun down (~1.5×10^9^ CFUs) and resuspended in a lysis mix containing 1x FastBreak lysis buffer, 3.8 mg/mL lysozyme, and ~25 units benzonase in instant ocean medium. Cells were allowed to lyse for 10 minutes at room temperature, and then mixed 1:1 with 2x SDS sample buffer [200 mM Tris pH 6.8, 25% glycerol, 1.8% SDS, 0.02% Bromophenol Blue, 5% β-mercaptoethanol]. Cell lysates were separated by SDS PAGE (10% gel) and then electrophoretically transferred to a PVDF membrane. This membrane was then blotted with primary mouse monoclonal α-FLAG M2 (Sigma) as described previously^64^ or with primary mouse monoclonal α-RpoA (Biolegend). Blots were washed and incubated with α-mouse secondary antibodies conjugated to horseradish peroxidase enzyme. Blots were developed with Pierce™ ECL Western Blotting Substrate and then imaged with a ProteinSimple FluorChem E system.

### Phylogenetic analysis

Phylogenetic tree of CdpA homologs among *Vibrionaceae* (**Fig. S6**) was constructed by first searching 287 publicly available, complete *Vibrio* genomes. A search with BUSCO v.4.0.5^65^ returned 214 HMMs that were complete, non-duplicated, and common to each assembly. The homologous gene sequences for these common HMMs from each assembly were identified and their corresponding CDS sequences were multiple aligned in the context of their corresponding amino acid sequences with MUSCLE v.3.8.31^66^. The multiple sequence alignments for individual genes were concatenated from which a best scoring maximum likelihood tree was generated with RAxML v.8.2.12^67^. We used “Interactive Tree Of Life” webserver v.6 (https://itol.embl.de) for visualizing the resulting tree. We identified homologs to *V. cholerae* CdpA among these 287 genomes and overlaid the corresponding percent identity values on the tree.

Phylogenetic tree based on cdpA homology observed among eubacterial genomes (**Fig. S7**) was generated by first searching the CdpA HMM profile built from the *V. cholerae* CdpA homologs among the *Vibrionales* against the eubacterial protein sequences (subset of NCBI nr database, taxonomy id = 2), using HMMER v.3.3.2 (http://hmmer.org/). Sequences for hits with a minimum alignment length of 700 amino acids were obtained and a multiple sequence alignment was performed on them using MUSCLE v.3.8.31^66^ (parameters: -diags -sv -distance1 kbit20_3). A best scoring maximum likelihood tree was generated using RAxML ver.8.2.12^67^. The nodes corresponding to the genus *Vibrio* were collapsed into a single node in blue, while those corresponding to the genus *Aliivibrio* were displayed in red.

The phylogenetic analysis showing presence or absence of CdpA homologs (**Fig. 4E**) was generated with a broadly representative set of genomes adapted from Wu and Eisen^68^. Phylosift^69^ identified and aligned 37 conserved protein sequences, which were subsequently concatenated into a combined alignment. RAxML^67^ estimated the maximum likelihood phylogeny using the LG amino acid substitution model with fixed residue frequencies and Gamma-distributed rate variation among sites.

Representative alignment of CdpA homologs of selected species was accomplished with T-Coffee, v.11.0^70,71^, and reformatted with BOXSHADE v.3.2.

### Detecting NO in liquid culture

NO production in liquid culture was determined with DAR 4M-AM, a fluorescent NO indicator dye. For measurements under conditions that induce NO production in *E. coli*, cells were first grown to an OD_600_ of ~2.5 at 30°C shaking in LB + 20 mM sodium nitrate. Cultures were then incubated statically at room temperature for 3 hr. Cultures were centrifuged and resuspended in either fresh LB (“-N”) or LB + 20 mM nitrate (“+N”) to a normalized OD_600_ of 2.5. DAR-4M AM was then added to each reaction to a final concentration of 12.5 μM.

For measurements under conditions that induce detachment in *V. cholerae* (“-air”), cells were first grown to an OD_600_ of ~2.5 at 30°C shaking in LB. 1 mL culture was loaded into a stoppered argon-flushed glass tube (microaerobic), DAR-4M AM was added to a final concentration of 12.5 μM, and the culture was then allowed to incubate at room temperature for 3 hr.

For measurements of NO production by a chemical NO donor, DEA-NONOate was added to final concentrations of 100 μM, 40 μM, or 10 μM in either fresh LB or cultures of *E. coli* grown in LB to an OD_600_ of ~2.5 at 30°C shaking (i.e., conditions where endogenous NO is not produced). DAR-4M AM was then added to each reaction to a final concentration of 12.5 μM.

Fluorescence of all samples was kinetically monitored with a BioTek Synergy H1M microplate reader with excitation set to 500 nm and emission set to 590 nm.

### CdpA purification and in vitro PDE activity assays

CdpA was cloned into a pET20b vector appending a 6xHis tag at the C terminus by the use of NdeI and XhoI restrictions enzymes. The resulting plasmid was transformed into BL21 (DE3) pLysS cells. Transformed cells were grown in yeast extract media (4.5% Yeast extract with 17 mM NaH_2_PO_4_ and 72 mM Na_2_HPO_4_, pH 7.5). Expression was induced at OD_595_ of 0.8 with 25 μM IPTG and carried out overnight at 16 °C. 0.1 mM ALA (δ-aminolevulinic acid) was added at the time of induction. For purification, cells were lysed by sonication in buffer containing 20 mM Tris-HCl, 250 mM KCl, 10% glycerol, 5 mM 2-mercaptoethanol, 50 μM EDTA, 1 mM PMSF, 1% Triton X-100, and 20 μM hemin, pH 8.0. Cleared lysate was loaded on a nickel-NTA (GE) column and the protein was eluted using an imidazole gradient. Three wash steps were performed: 100 ml of 10 mM imidazole, 50 ml of 20 mM imidazole and 5 ml of 50 mM imidazole. After washing, the proteins were eluted in buffer with 250 mM imidazole and desalted with a PD-10 column (GE) in buffer containing 50 mM Tris, 250 mM KCl, 5 mM 2-mercaptoethanol, pH 8.0.

PDE assays were performed at 25 °C overnight in an assay buffer containing 50 mM Tris-HCl, 5 mM MgCl_2_, 50 μM c-di-GMP and pH 8.0. Reactions were initiated by the addition of 50 nM of purified protein. Reactions were quenched by addition of 10 mM CaCl_2_ and subsequent heating to 95 °C for 5 min to precipitate protein. Precipitated protein was removed by centrifugation and the resulting supernatant was filtered through a 0.22 μm membrane and analyzed by HPLC on a reverse phase C18 column (Shimazu) with an ion pairing buffer system.

### ΔNap growth assay

Cells were inoculated into M9 + 1% glucose +/- 20 mM sodium nitrate in a stoppered argon-flushed glass tube (microaerobic) and allowed to grow for 24 hr at 30°C static. Growth was assessed by measuring the final OD_600_ of cultures.

### Statistics

Statistical differences were determined using one-way ANOVA tests and post-hoc Holm-Šídák test, α = 0.5, or by two-tailed Student’s t-test, as noted in the figure legends. Statistical comparisons for MixD, binding frequency, western blot, piliation, and initial binding experiments were performed on the log-transformed data. All statistical comparisons were performed with GraphPad Prism software version 9.0.1. All means and statistical comparisons can be found in **Supplemental Tables 4** and **5**, respectively.

### Data availability

The data that support the findings of this study are available from the corresponding author upon request.

## Supplemental Information

**Supplemental Figure 1:**
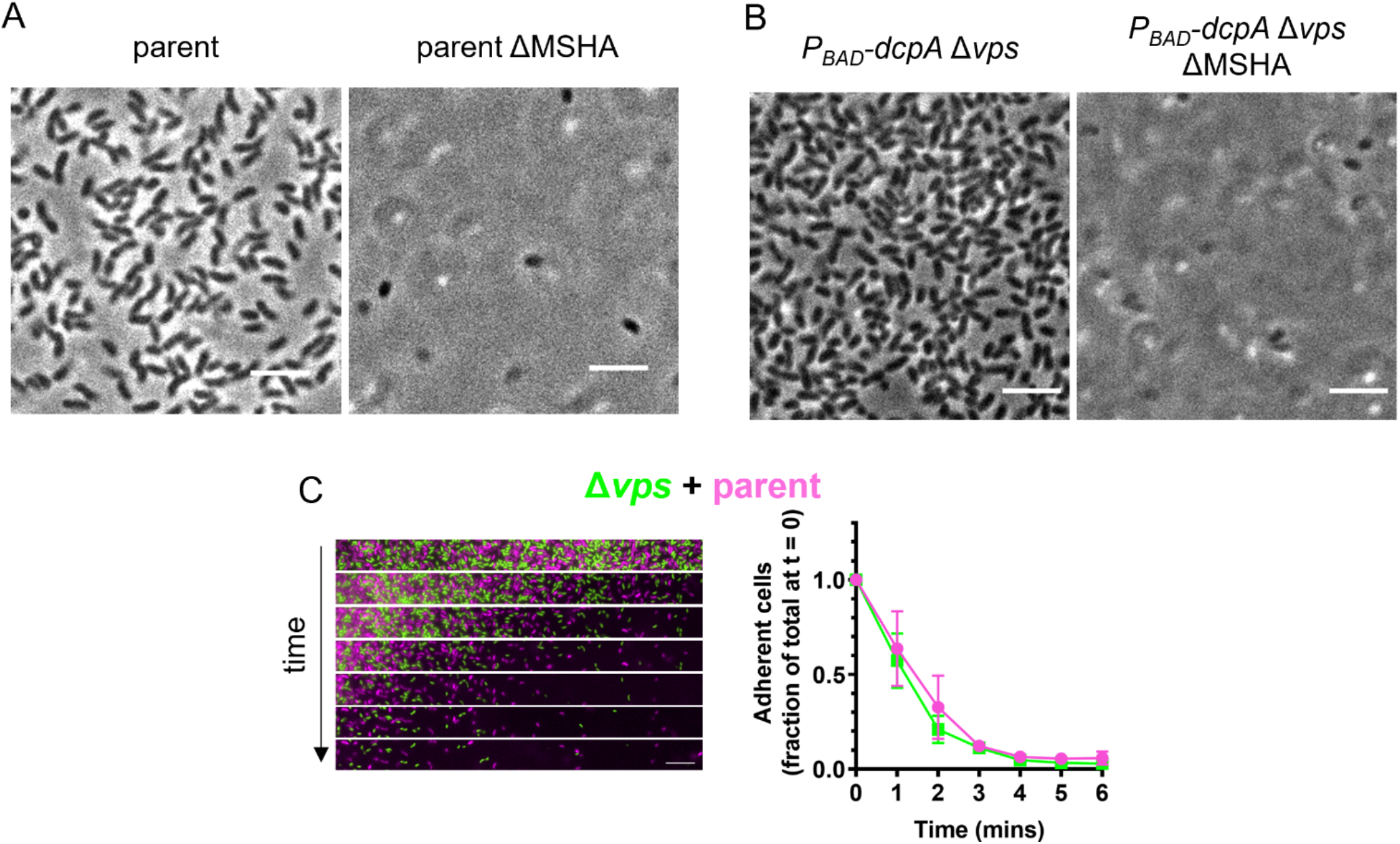
MSHA pili, but not *vps*, are required for initial cell attachment. (**A**-**B**) Representative phase contrast images showing surface binding in the indicated strain backgrounds. ΔMSHA signifies deletion of the entire MSHA pilus operon (ΔVC0399 – VC0414). Importantly, deleting the MSHA pilus operon from the P_BAD_-*dcpA* Δ*vps* strain prevented attachment, even under induction of *dcpA* (cells in **B** were grown with 0.15% arabinose), indicating that attachment is dependent on MSHA pili in this background. Scale bar = 5 μm. (**C**) MixD assay of Δ*vps* expressing GFP (green) and a parent strain expressing mCherry (fuchsia). Representative montage of time-lapse imaging (left) with 1-minute intervals between frames. Scale bar = 10 μm. Quantification of three biological replicates is shown in the line graph (right) and is displayed as the mean ± SD.

**Supplemental Figure 2:**
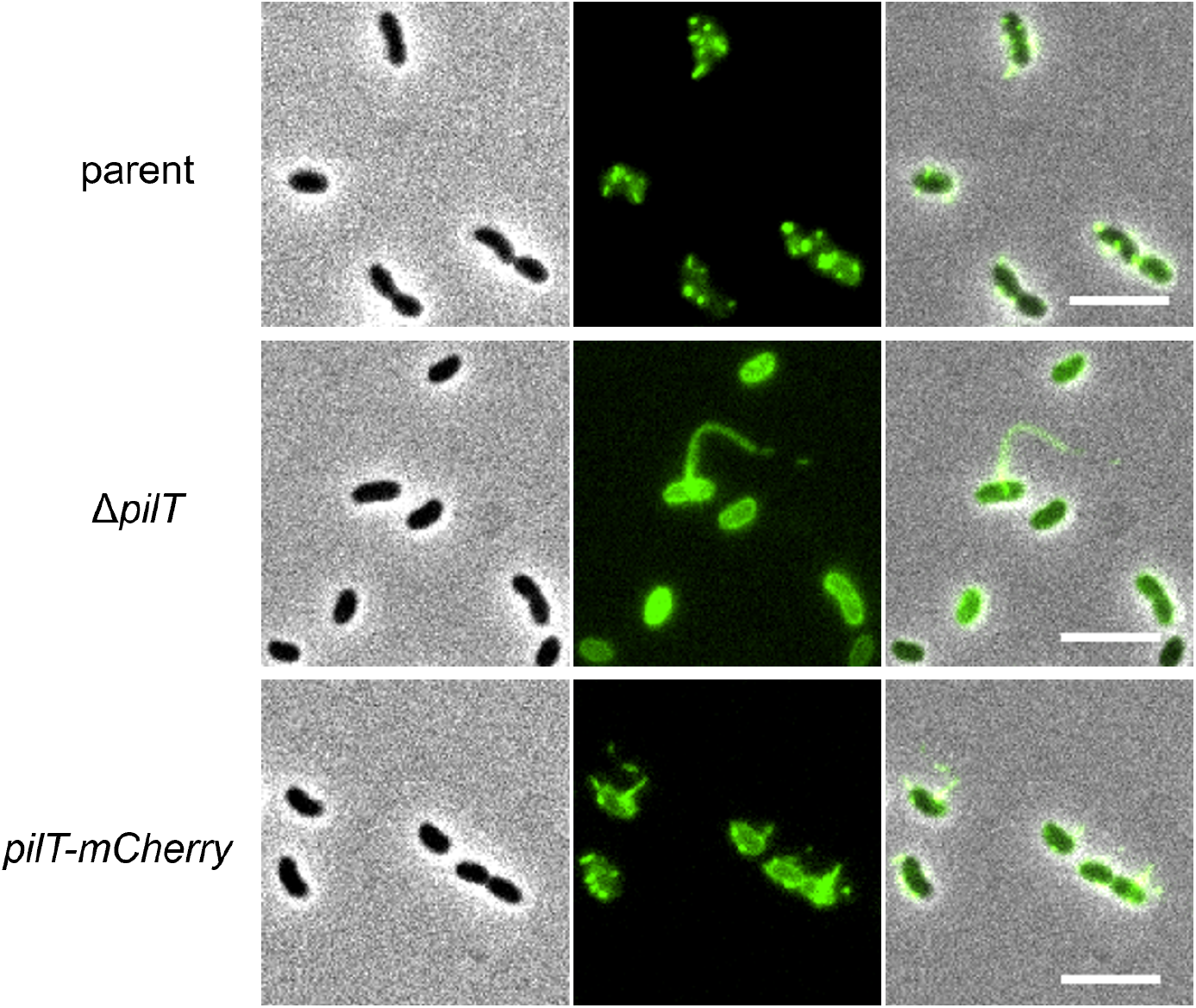
*PilT-mCherry* cells retain MSHA surface piliation, unlike Δ*pilT* cells. Representative images showing piliation states of parent, Δ*pilT*, and *pilT-mCherry* strains. Phase images (left) show cell boundaries, FITC images (middle) show AF488-mal labelled MSHA pili, and the overlay is shown on the right. Production of pili is rare in Δ*pilT* cells, but when observed, cells make a single long pilus as depicted in the representative image. Scale bar = 5 μm.

**Supplemental Figure 3:**
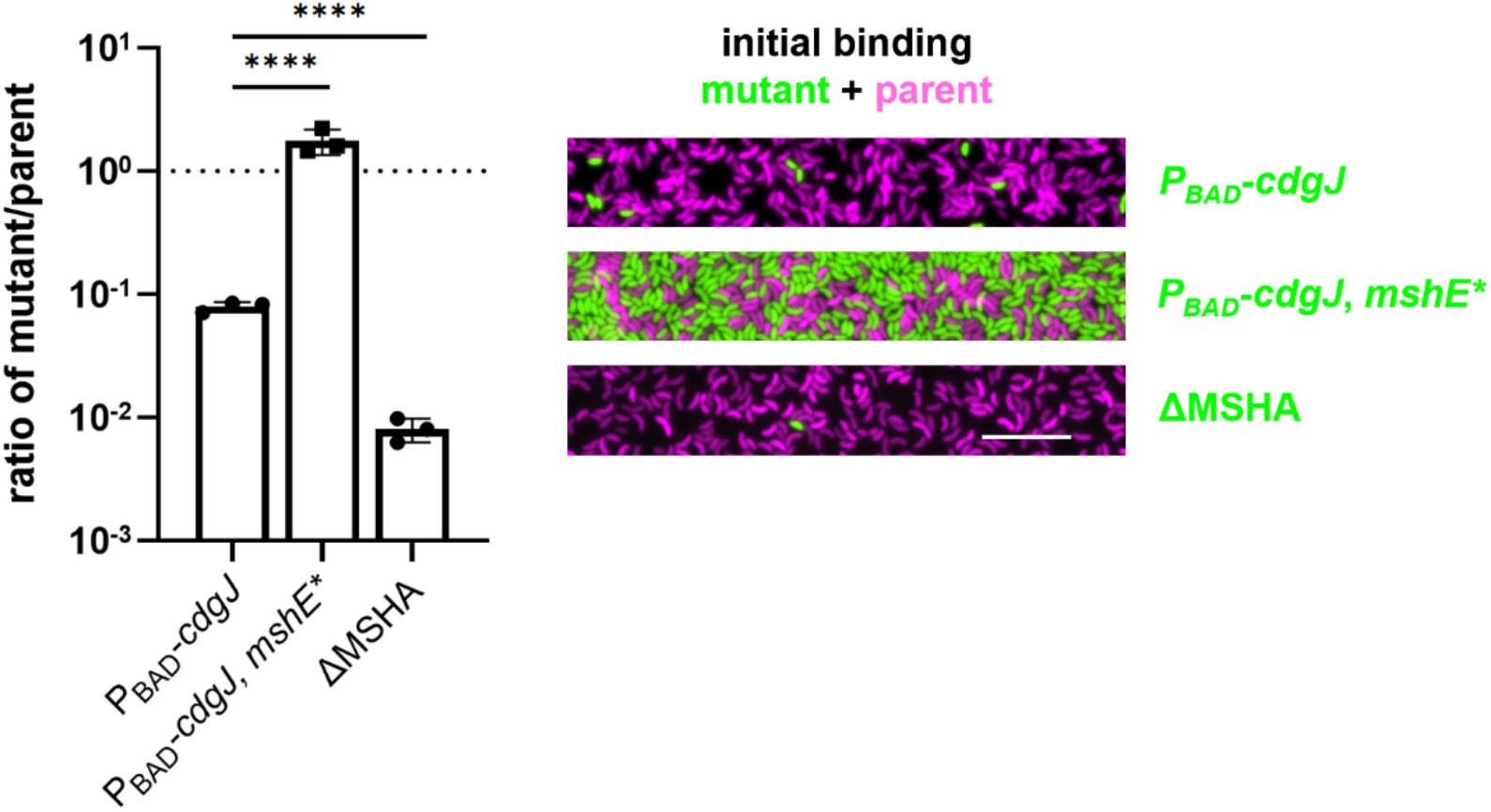
Increasing PDE activity prevents cell attachment in a manner that is dependent on the allosteric regulation of MshE activity. Quantification (left) of initial attachment of a 1:1 mixture of the parent and the mutant strain indicated on the X-axis. Ratio is determined by dividing the number of attached mCherry expressing parent cells (fuchsia) by the number of attached GFP-expressing mutant cells (green) as depicted in the representative images shown on the right. If strains bind equally well, the expected ratio is 1, which is indicated by the dotted line. Data are from three independent experiments and shown as the mean ± SD with at least 240 cells analyzed per replicate. Statistical comparisons were made with ANOVA and post-hoc Holm-Šídák test. \*\*\*\**P* < 0.0001. Scale bar = 10 μm.

**Supplemental Figure 4:**
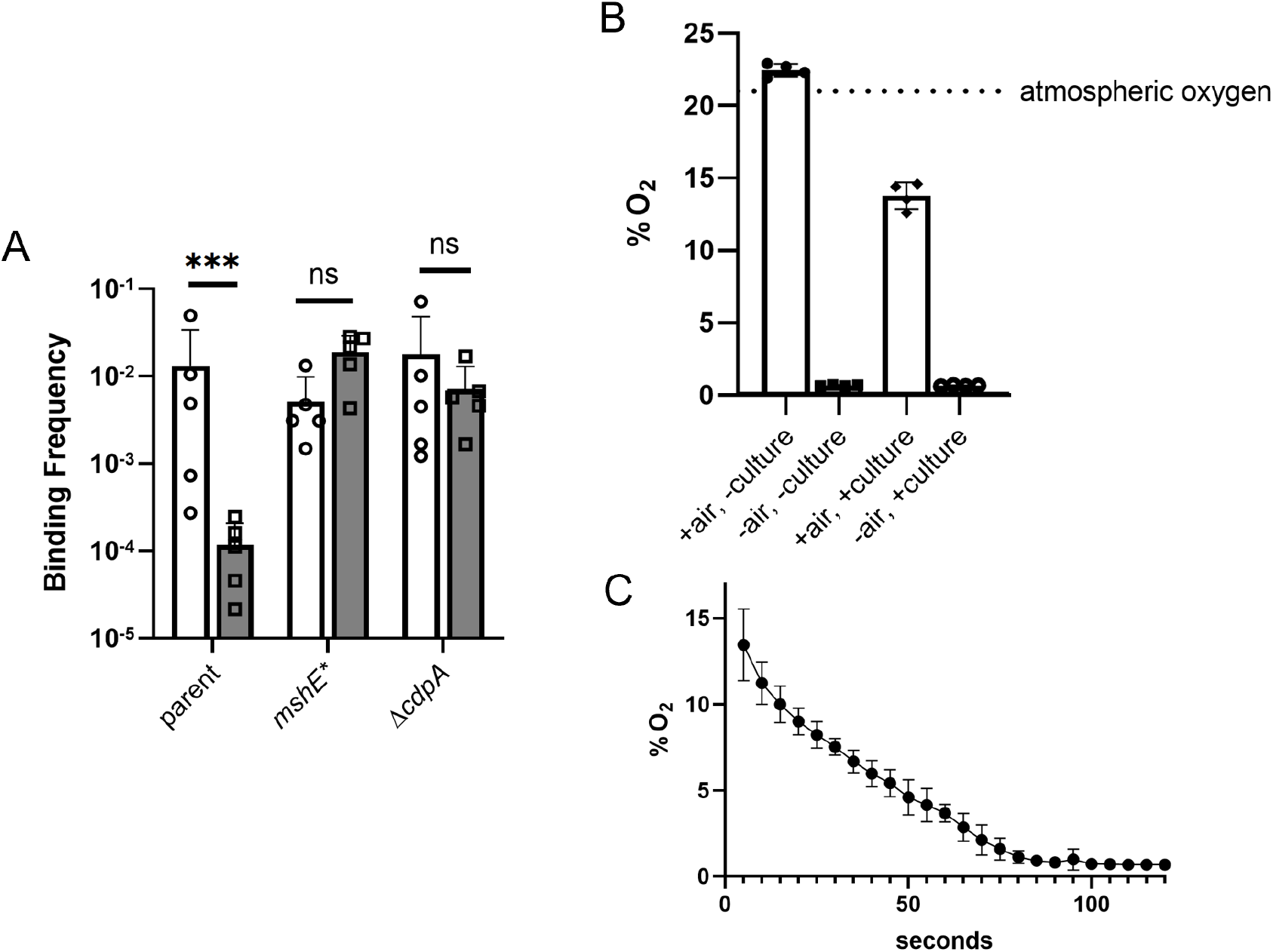
Large-scale binding assays in glass culture tubes match MixD phenotypes. (**A**) Large-scale binding assays of cultures incubated in aerobic (open bars) or microaerobic conditions (shaded bars). Data are from five independent biological replicates and shown as the mean ± SD. Statistical comparisons were made by one-way ANOVA and post-hoc Holm-Šídák test. \*\*\**P* < 0.001. (**B**) Oxygen levels in aerobic vs. microaerobic growth conditions. “+ air” indicates tubes that were incubated rolling, open to the air. “-air” indicates tubes that were flushed with argon and sealed. Initial % O_2_ is displayed in the open bars, and % O_2_ after incubation with culture is displayed in the shaded bars. (**C**) Oxygen levels of the aerobic culture upon transition from rolling to static incubation. Graphs **B** and **C** represent 4 independent biological replicates and are shown as the mean ± SD.

**Supplemental Figure 5:**
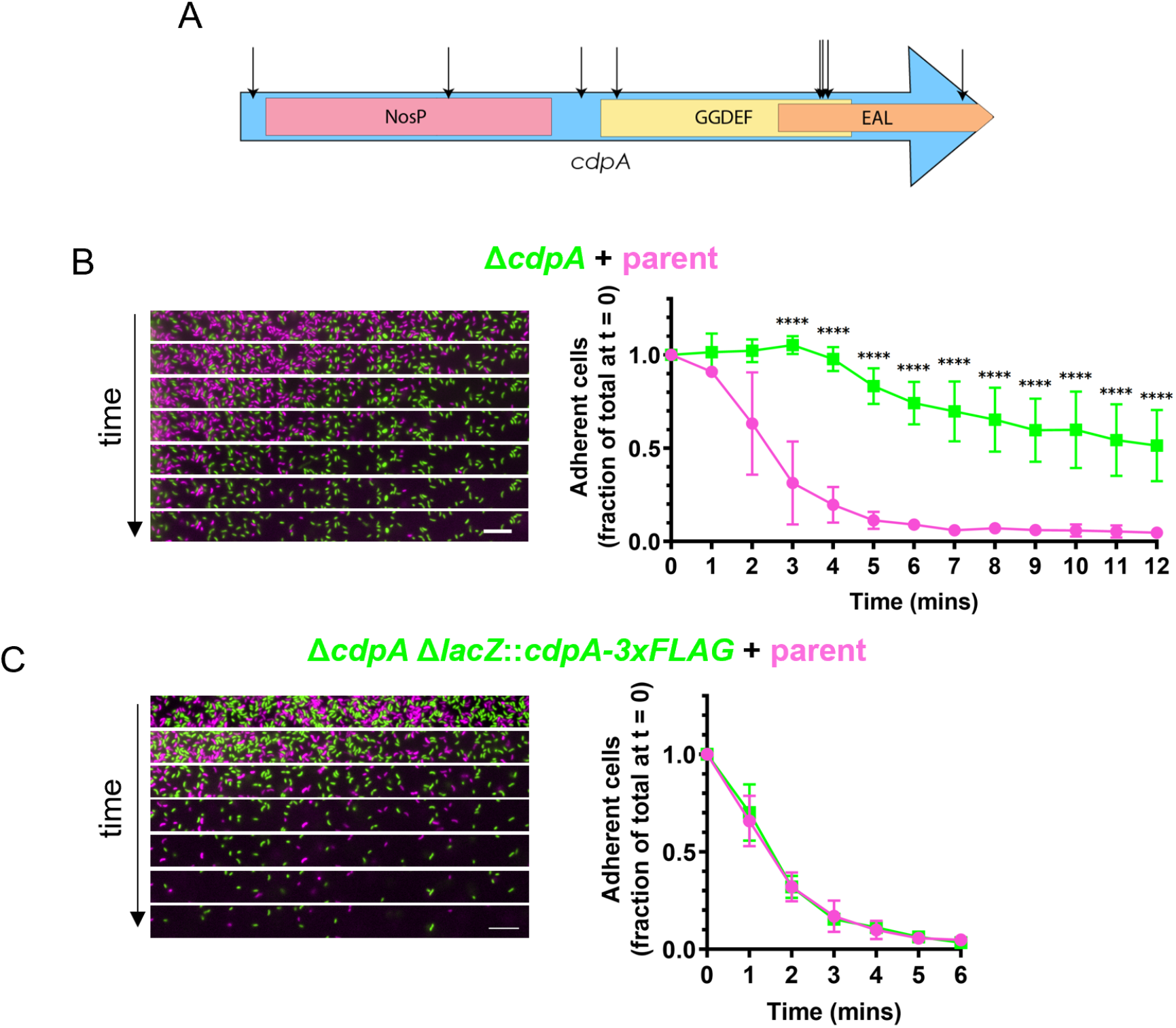
CdpA is required for optimal cell detachment from slides. (**A**) Schematic of the domain architecture of CdpA. Black arrows indicate transposon insertion sites recovered during our genetic selection. (**B**-**C**) MixD assays of Δ*cdpA* and *cdpA-3xFLAG* strains expressing GFP (green) with a parent strain expressing mCherry (fuschia). Representative montages of time-lapse imaging (left) with 1-minute intervals between frames. Scale bar = 10 μm. Quantification of three biological replicates is shown in each line graph (right) and is displayed as the mean ± SD. Statistical comparisons were made by one-way ANOVA and post-hoc Holm-Šídák test. \*\*\*\**P* < 0.0001.

**Supplemental Figure 6:**
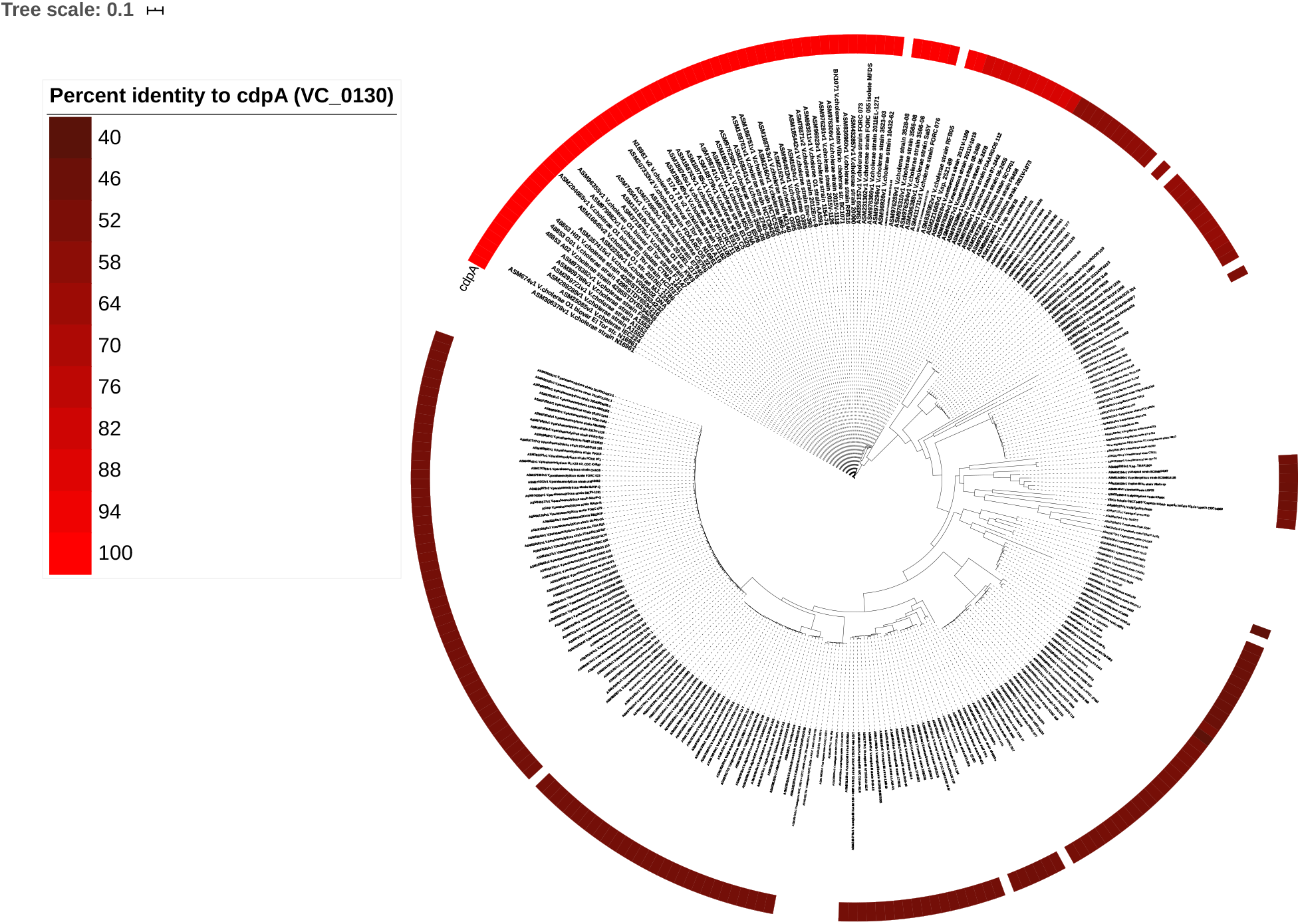
CdpA is widely conserved among the *Vibrionaceae*. Phylogenetic tree of *Vibrio* genus derived from a multi-locus protein multiple sequence alignment. Strains that encode homologs of *cdpA* are denoted with a red bar, and the brightness of the bar corresponds to the percent identity with *V. cholerae* CdpA (VC0130).

**Supplemental Figure 7:**
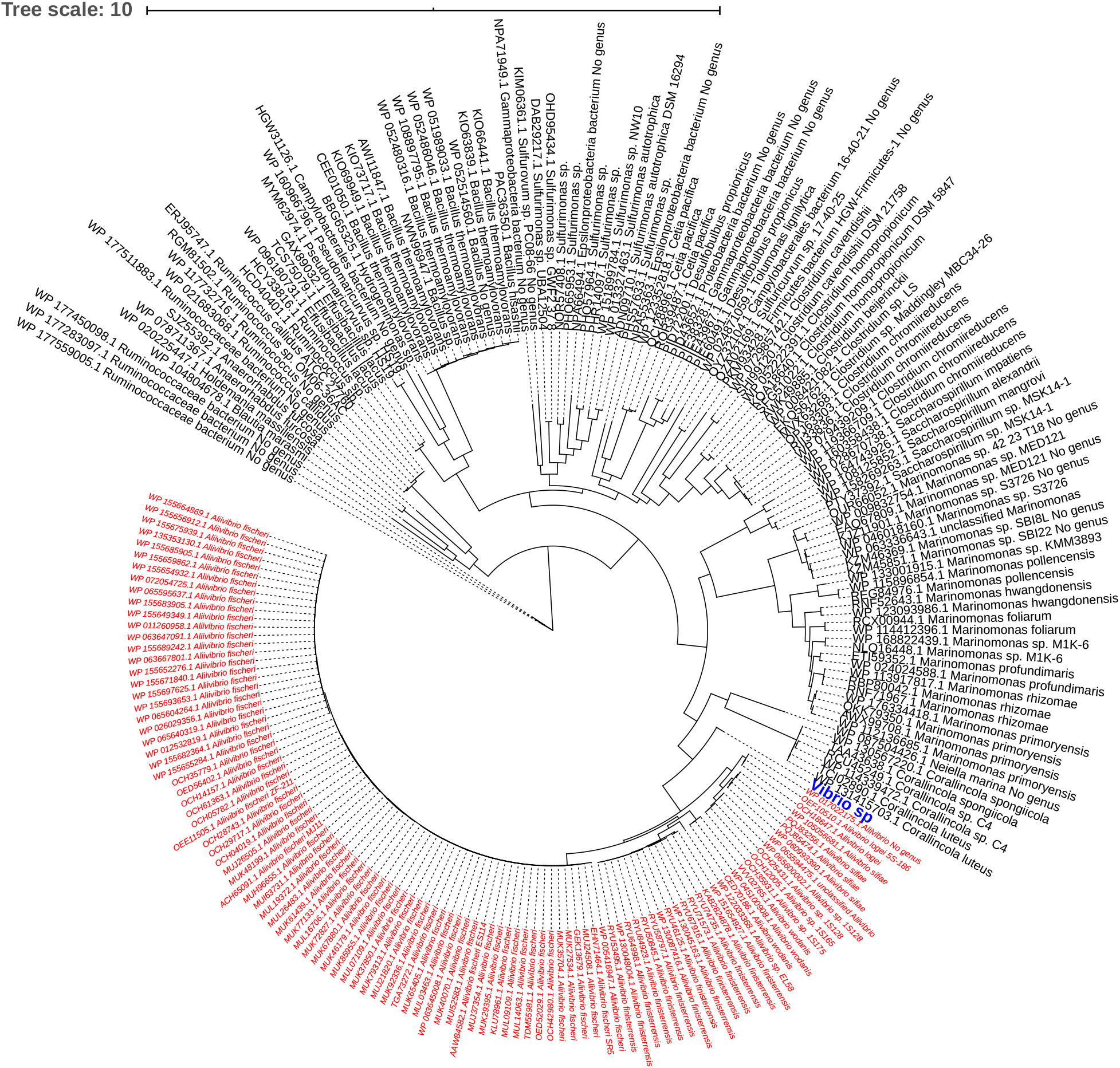
CdpA homologs are found in diverse eubacteria. Phylogenetic tree of CdpA containing eubacteria derived from a CpdA protein multiple sequence alignment. *Vibrio* species (blue) are condensed into a single node and *Allivibrio* species are designated by red text.

**Supplemental Figure 8:**
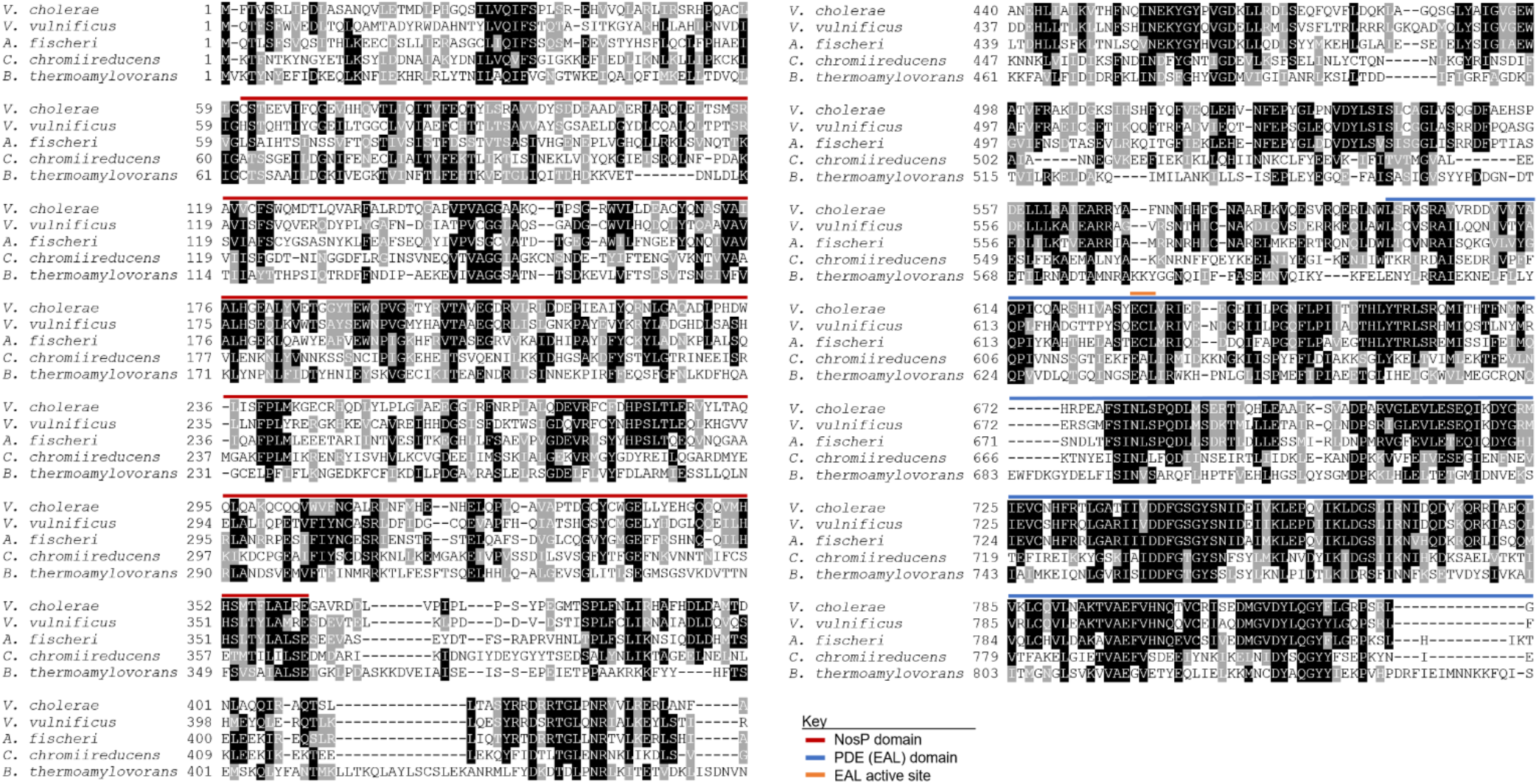
Alignment of CdpA homologs from different species. Multiple sequence alignment of CdpA homologs identified from *V. cholerae, Vibrio vulnificus, Allivibrio fischeri, Clostridium chromiireducens*, and *Bacillus thremoamylovorans*. The red line above the sequences denotes the NosP domain. The blue line above the sequences indicates the EAL domain responsible for PDE activity. The orange line marks the residues that give the EAL domain its name (ECL in *Vc* CdpA). Residues highlighted in black are identical, while those highlighted in gray are similar. The GenBank Accession numbers used for the protein sequences are as follows: *V. cholerae* (QJS76791.1), *V. vulnificus* (WP_131070112.1), *A. fischeri* (WP_155689242.1), *C. chromiireducens* (WP_079439209.1), *B. thermoamylovorans* (WP_108897795.1).

**Supplemental Figure 9:**
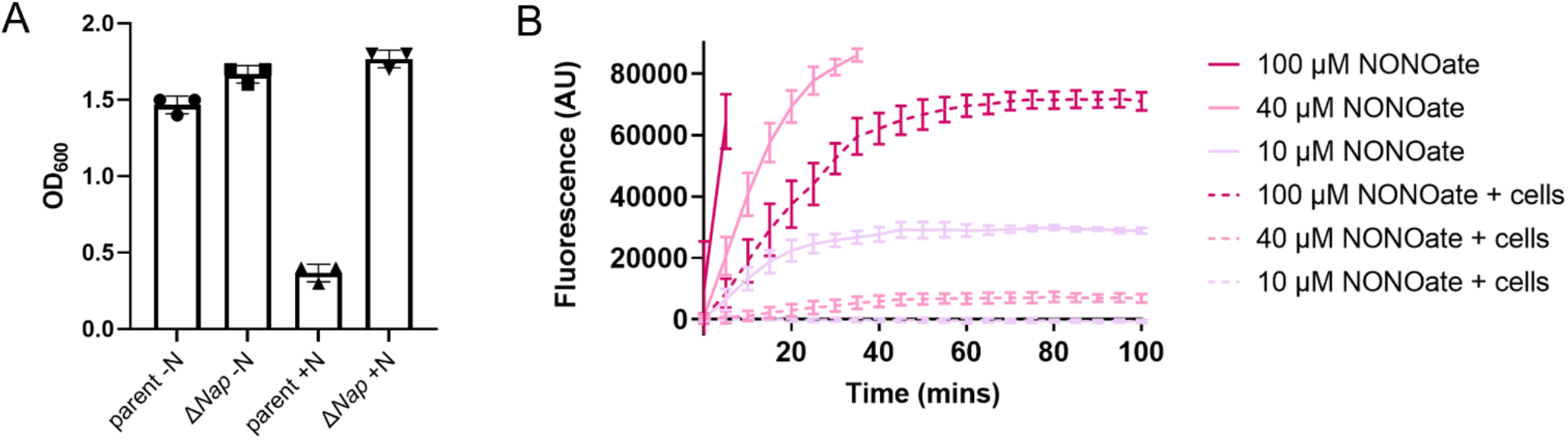
Validation of Δ*Nap* and DAR 4M-AM assays. (**A**) Microaerobic growth of parent and Δ*Nap* after 24hr in M9 + 1% glucose either with (+N) or without (-N) 20mM nitrate. These results demonstrate that the parent grows poorly with added nitrate compared to the Δ*Nap* mutant, which is consistent with elevated levels of nitrite inhibiting bacterial growth in the parent as previously described^39^. This confirms that our Δ*Nap* mutant behaves as expected. (**B**) DAR 4M-AM measurement of NO released by DEA-NONOate either with (dashed lines) or without (solid lines) live bacterial cells present. Quantification of three biological replicates is shown in each graph and is displayed as the mean ± SD.

**Supplemental Movie 1: MSHA pilus retraction precedes detachment.**

Pilus retraction captured by epifluorescence time-lapse microscopy of AF488-mal labeled cells, as in **Fig. 1A**. The capture interval is 5 s between each frame. Scale bar = 1 μm.

**Supplemental Movie 2: Parent cells detach during MixD analysis.**

Time lapse of MixD assays of parent cells expressing either GFP (green) or mCherry (fuchsia), as in **Fig. 1C**. The capture interval is 1 min between each frame. Scale bar = 20 μm.

**Supplemental Movie 3: PilT-mCherry cells stay attached during MixD analysis**

Time lapse of MixD assays of PilT-mCherry cells expressing GFP (green) or parent cells expressing mCherry (fuchsia), as in **Fig. 1D**. The capture interval is 1 min between each frame. Scale bar = 20 μm.

**Supplemental Movie 4: PilT-mCherry attachment correlates with maintained pilus extension.**

Time lapse of pilus retraction captured by epifluorescence microscopy of AF488-mal labeled cells, as portrayed in **Fig. 1E**. Parent cells are distinguished by CyPet expression (blue). The capture interval is 5 s between each frame. Scale bar = 1 μm.

**Supplemental Movie 5: Aerobic GFP production only occurs at the edge of wells.**

Time lapse of a mixed culture of parent cells expressing mCherry (fuchsia) with *mshE*\* cells expressing CyPet (blue). The *mshE*\* also contains a P_BAD_-*gfp* construct (top), which serves as a bioreporter for O_2_ in this experiment. Arabinose was added to the mixed culture immediately before imaging, as in **Fig. 3B**. The capture interval is 5 mins between each frame. Scale bar = 20 μm.

**Supplemental Table 1: CdpA homologs found in eubacterial species.**

See attached file.

**Supplemental Table 2:**
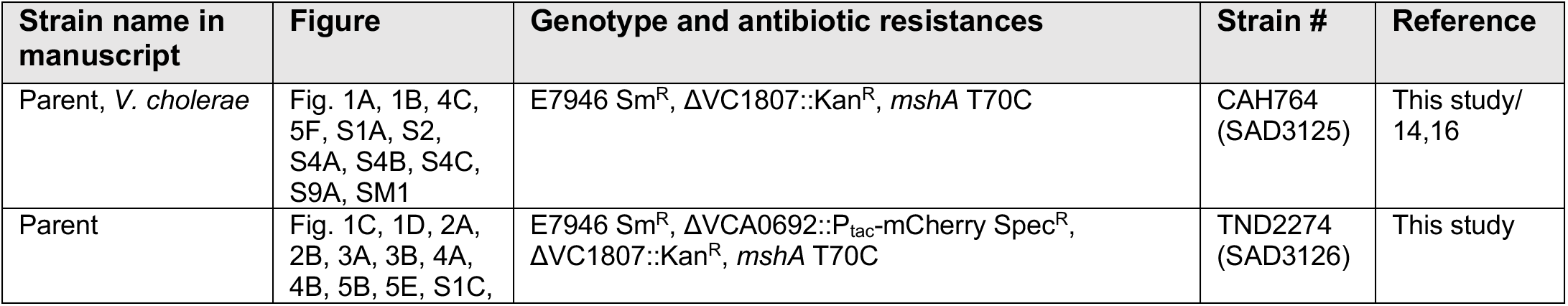
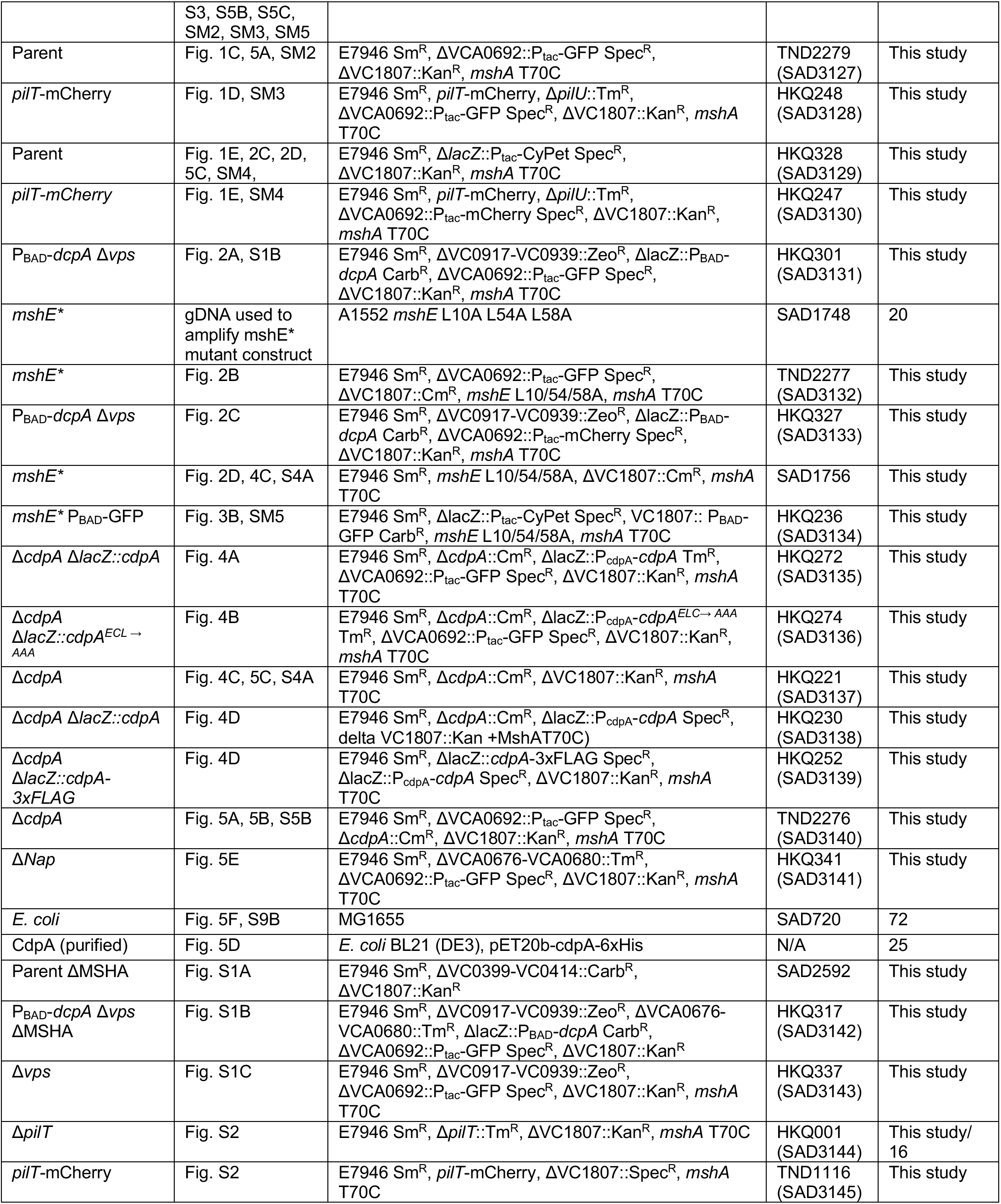
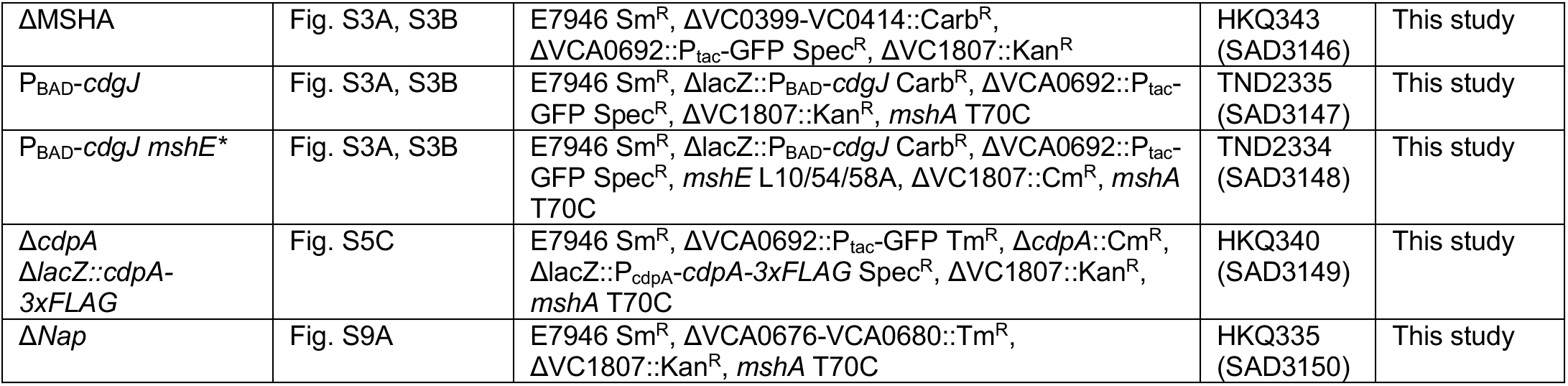
Strain List

**Supplemental Table 3:**
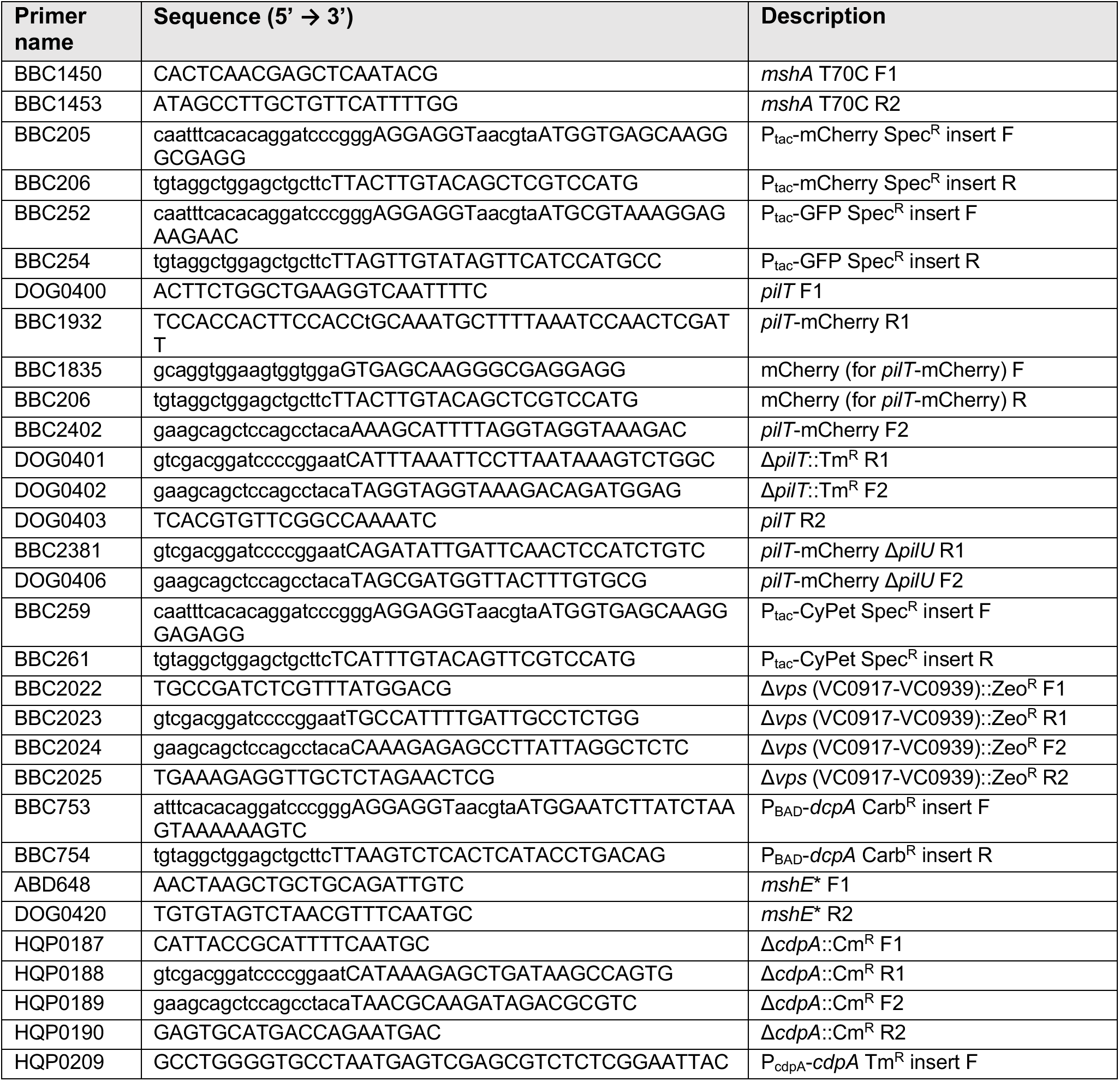
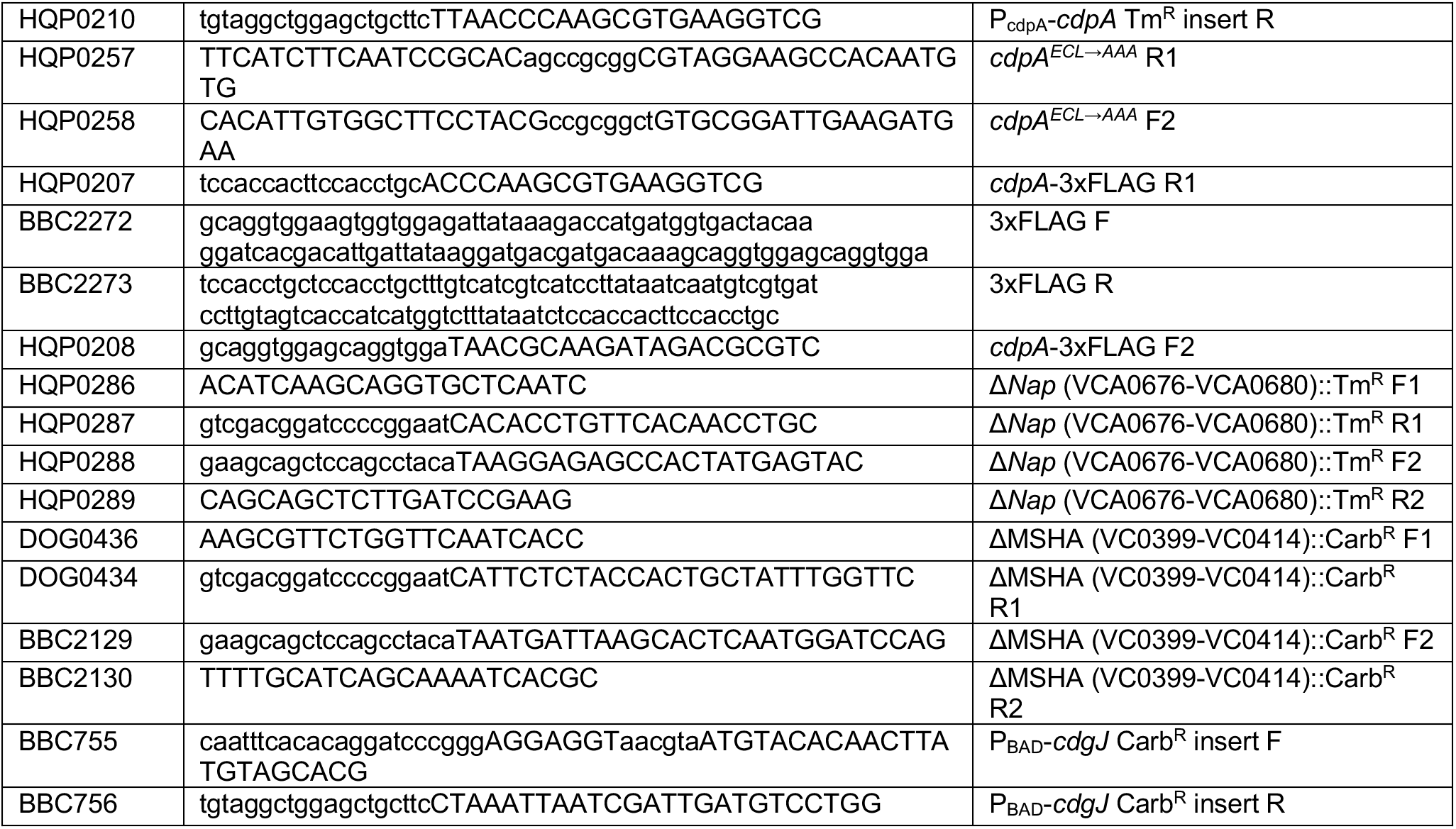
Primer List

**Supplemental Table 4:** Mean values for each data set

| Figure | Strain name | Mean | Figure | Strain name | Mean |
| --- | --- | --- | --- | --- | --- |
| Fig. 1C | Parent (green), time = 0 | 1.00E+00 | Fig. 5E | ΔNap, time = 2 | 2.43E-01 |
| Fig. 1C | Parent (green), time = 1 | 6.59E-01 | Fig. 5E | ΔNap, time = 3 | 1.39E-01 |
| Fig. 1C | Parent (green), time = 2 | 3.42E-01 | Fig. 5E | ΔNap, time = 4 | 8.95E-02 |
| Fig. 1C | Parent (green), time = 3 | 1.99E-01 | Fig. 5E | ΔNap, time = 5 | 7.16E-02 |
| Fig. 1C | Parent (green), time = 4 | 1.30E-01 | Fig. 5E | ΔNap, time = 6 | 5.00E-02 |
| Fig. 1C | Parent (green), time = 5 | 1.24E-01 | Fig. 5E | Parent, time = 0 | 1.00E+00 |
| Fig. 1C | Parent (green), time = 6 | 8.96E-02 | Fig. 5E | Parent, time = 1 | 7.34E-01 |
| Fig. 1C | Parent (fuchsia), time = 0 | 1.00E+00 | Fig. 5E | Parent, time = 2 | 3.31E-01 |
| Fig. 1C | Parent (fuchsia), time = 1 | 6.14E-01 | Fig. 5E | Parent, time = 3 | 1.92E-01 |
| Fig. 1C | Parent (fuchsia), time = 2 | 2.69E-01 | Fig. 5E | Parent, time = 4 | 1.00E-01 |
| Fig. 1C | Parent (fuchsia), time = 3 | 1.22E-01 | Fig. 5E | Parent, time = 5 | 7.24E-02 |
| Fig. 1C | Parent (fuchsia), time = 4 | 8.78E-02 | Fig. 5E | Parent, time = 6 | 6.66E-02 |
| Fig. 1C | Parent (fuchsia), time = 5 | 7.74E-02 | Fig. S1C | Δvps, time = 0 | 1.00E+00 |
| Fig. 1C | Parent (fuchsia), time = 6 | 6.29E-02 | Fig. S1C | Δvps, time = 1 | 5.72E-01 |
| Fig. 1D | *pilT*-mCherry, time = 0 | 1.00E+00 | Fig. S1C | Δvps, time = 2 | 2.10E-01 |
| Fig. 1D | *pilT*-mCherry, time = 1 | 1.04E+00 | Fig. S1C | Δvps, time = 3 | 1.13E-01 |
| Fig. 1D | *pilT*-mCherry, time = 2 | 1.06E+00 | Fig. S1C | Δvps, time = 4 | 4.75E-02 |
| Fig. 1D | *pilT*-mCherry, time = 3 | 1.07E+00 | Fig. S1C | Δvps, time = 5 | 3.35E-02 |
| Fig. 1D | *pilT*-mCherry, time = 4 | 1.09E+00 | Fig. S1C | Δvps, time = 6 | 2.83E-02 |
| Fig. 1D | *pilT*-mCherry, time = 5 | 1.10E+00 | Fig. S1C | Parent, time = 0 | 1.00E+00 |
| Fig. 1D | *pilT*-mCherry, time = 6 | 1.09E+00 | Fig. S1C | Parent, time = 1 | 6.37E-01 |
| Fig. 1D | Parent, time = 0 | 1.00E+00 | Fig. S1C | Parent, time = 2 | 3.26E-01 |
| Fig. 1D | Parent, time = 1 | 7.26E-01 | Fig. S1C | Parent, time = 3 | 1.21E-01 |
| Fig. 1D | Parent, time = 2 | 4.64E-01 | Fig. S1C | Parent, time = 4 | 6.42E-02 |
| Fig. 1D | Parent, time = 3 | 2.78E-01 | Fig. S1C | Parent, time = 5 | 5.51E-02 |
| Fig. 1D | Parent, time = 4 | 1.93E-01 | Fig. S1C | Parent, time = 6 | 5.80E-02 |
| Fig. 1D | Parent, time = 5 | 1.25E-01 | Fig. S3 | $P_{BAD-cdgJ}$ | 7.95E-02 |
| Fig. 1D | Parent, time = 6 | 9.31E-02 | Fig. S3 | $P_{BAD-cdgJ}, mshE^*$ | 1.76E+00 |
| Fig. 2A | $P_{BAD-dcpA} \Delta vps$, time = 0 | 1.00E+00 | Fig. S3 | $\Delta MSHA$ | 8.05E-03 |
| Fig. 2A | $P_{BAD-dcpA} \Delta vps$, time = 1 | 1.18E+00 | Fig. S4A | Parent, aerobic | 1.32E-02 |
| Fig. 2A | $P_{BAD-dcpA} \Delta vps$, time = 2 | 1.29E+00 | Fig. S4A | Parent, microaerobic | 1.17E-04 |
| Fig. 2A | $P_{BAD-dcpA} \Delta vps$, time = 3 | 1.36E+00 | Fig. S4A | MshE*, aerobic | 5.12E-03 |
| Fig. 2A | $P_{BAD-dcpA} \Delta vps$, time = 4 | 1.40E+00 | Fig. S4A | MshE*, microaerobic | 1.89E-02 |
| Fig. 2A | $P_{BAD-dcpA} \Delta vps$, time = 5 | 1.45E+00 | Fig. S4A | $\Delta cdpA$, aerobic | 1.78E-02 |
| Fig. 2A | $P_{BAD-dcpA} \Delta vps$, time = 6 | 1.48E+00 | Fig. S4A | $\Delta cdpA$, microaerobic | 7.12E-03 |
| Fig. 2A | Parent, time = 0 | 1.00E+00 | Fig. S4B | + air, - culture | 2.25E+01 |
| Fig. 2A | Parent, time = 1 | 7.49E-01 | Fig. S4B | - air, - culture | 6.73E-01 |
| Fig. 2A | Parent, time = 2 | 3.28E-01 | Fig. S4B | + air, + culture | 1.38E+01 |
| Fig. 2A | Parent, time = 3 | 1.58E-01 | Fig. S4B | - air, + culture | 6.55E-01 |
| Fig. 2A | Parent, time = 4 | 8.90E-02 | Fig. S4C | 5 seconds | 1.35E+01 |
| Fig. 2A | Parent, time = 5 | 5.34E-02 | Fig. S4C | 10 seconds | 1.12E+01 |
| Fig. 2A | Parent, time = 6 | 6.41E-02 | Fig. S4C | 15 seconds | 1.00E+01 |
| Fig. 2B | $mshE^*$, time = 0 | 1.00E+00 | Fig. S4C | 20 seconds | 9.00E+00 |
| Fig. 2B | $mshE^*$, time = 1 | 1.12E+00 | Fig. S4C | 25 seconds | 8.23E+00 |
| Fig. 2B | $mshE^*$, time = 2 | 1.13E+00 | Fig. S4C | 30 seconds | 7.53E+00 |
| Fig. 2B | $mshE^*$, time = 3 | 1.10E+00 | Fig. S4C | 35 seconds | 6.68E+00 |
| Fig. 2B | $mshE^*$, time = 4 | 1.08E+00 | Fig. S4C | 40 seconds | 5.98E+00 |
| Fig. 2B | $mshE^*$, time = 5 | 1.12E+00 | Fig. S4C | 45 seconds | 5.43E+00 |
| Fig. 2B | $mshE^*$, time = 6 | 1.14E+00 | Fig. S4C | 50 seconds | 4.60E+00 |
| Fig. 2B | Parent, time = 0 | 1.00E+00 | Fig. S4C | 55 seconds | 4.15E+00 |
| Fig. 2B | Parent, time = 1 | 8.82E-01 | Fig. S4C | 60 seconds | 3.68E+00 |
| Fig. 2B | Parent, time = 2 | 5.76E-01 | Fig. S4C | 65 seconds | 2.85E+00 |
| Fig. 2B | Parent, time = 3 | 3.41E-01 | Fig. S4C | 70 seconds | 2.12E+00 |
| Fig. 2B | Parent, time = 4 | 2.38E-01 | Fig. S4C | 75 seconds | 1.59E+00 |
| Fig. 2B | Parent, time = 5 | 1.65E-01 | Fig. S4C | 80 seconds | 1.13E+00 |
| Fig. 2B | Parent, time = 6 | 9.67E-02 | Fig. S4C | 85 seconds | 9.15E-01 |
| Fig. 4A | $\Delta cdpA, \Delta lacZ::cdpA$, time = 0 | 1.00E+00 | Fig. S4C | 90 seconds | 8.15E-01 |
| Fig. 4A | $\Delta cdpA, \Delta lacZ::cdpA$, time = 1 | 7.73E-01 | Fig. S4C | 95 seconds | 9.85E-01 |
| Fig. 4A | $\Delta cdpA, \Delta lacZ::cdpA$, time = 2 | 3.37E-01 | Fig. S4C | 100 seconds | 7.33E-01 |
| Fig. 4A | $\Delta cdpA, \Delta lacZ::cdpA$, time = 3 | 1.76E-01 | Fig. S4C | 105 seconds | 7.13E-01 |
| Fig. 4A | $\Delta cdpA, \Delta lacZ::cdpA$, time = 4 | 1.01E-01 | Fig. S4C | 110 seconds | 6.85E-01 |
| Fig. 4A | $\Delta cdpA, \Delta lacZ::cdpA$, time = 5 | 6.91E-02 | Fig. S4C | 115 seconds | 6.90E-01 |
| Fig. 4A | $\Delta cdpA, \Delta lacZ::cdpA$, time = 6 | 8.21E-02 | Fig. S4C | 120 seconds | 6.80E-0 |
| Fig. 4A | Parent, time = 0 | 1.00E+00 | Fig. S5B | $\Delta cdpA$, time = 0 | 1.00E+00 |
| Fig. 4A | Parent, time = 1 | 7.08E-01 | Fig. S5B | $\Delta cdpA$, time = 1 | 1.01E+00 |
| Fig. 4A | Parent, time = 2 | 2.59E-01 | Fig. S5B | $\Delta cdpA$, time = 2 | 1.02E+00 |
| Fig. 4A | Parent, time = 3 | 1.11E-01 | Fig. S5B | $\Delta cdpA$, time = 3 | 1.05E+00 |
| Fig. 4A | Parent, time = 4 | 7.49E-01 | Fig. S5B | $\Delta cdpA$, time = 4 | 9.87E-01 |
| Fig. 4A | Parent, time = 5 | 6.36E-01 | Fig. S5B | $\Delta cdpA$, time = 5 | 8.33E-01 |
| Fig. 4A | Parent, time = 6 | 3.77E-01 | Fig. S5B | $\Delta cdpA$, time = 6 | 7.41E-01 |
| Fig. 4B | $\Delta cdpA, \Delta lacZ::cdpA^{ECL \rightarrow AAA}$, time = 0 | 1.00E+00 | Fig. S5B | $\Delta cdpA$, time = 7 | 6.97E-01 |
| Fig. 4B | $\Delta cdpA, \Delta lacZ::cdpA^{ECL \rightarrow AAA}$, time = 1 | 1.10E+00 | Fig. S5B | $\Delta cdpA$, time = 8 | 6.52E-01 |
| Fig. 4B | $\Delta cdpA, \Delta lacZ:: cdpA^{ECL \rightarrow AAA}$, time = 2 | 1.14E+00 | Fig. S5B | $\Delta cdpA$, time = 9 | 5.96E-01 |
| Fig. 4B | $\Delta cdpA, \Delta lacZ:: cdpA^{ECL \rightarrow AAA}$, time = 3 | 1.16E+00 | Fig. S5B | $\Delta cdpA$, time = 10 | 5.98E-01 |
| Fig. 4B | $\Delta cdpA, \Delta lacZ:: cdpA^{ECL \rightarrow AAA}$, time = 4 | 1.09E+00 | Fig. S5B | $\Delta cdpA$, time = 11 | 5.43E-01 |
| Fig. 4B | $\Delta cdpA, \Delta lacZ:: cdpA^{ECL \rightarrow AAA}$, time = 5 | 1.05E+00 | Fig. S5B | $\Delta cdpA$, time = 12 | 5.14E-01 |
| Fig. 4B | $\Delta cdpA, \Delta lacZ:: cdpA^{ECL \rightarrow AAA}$, time = 6 | 1.03E+00 | Fig. S5B | Parent, time = 0 | 1.00E+00 |
| Fig. 4B | Parent, time = 0 | 1.00E+00 | Fig. S5B | Parent, time = 1 | 9.10E-01 |
| Fig. 4B | Parent, time = 1 | 6.75E-01 | Fig. S5B | Parent, time = 2 | 6.32E-01 |
| Fig. 4B | Parent, time = 2 | 3.84E-01 | Fig. S5B | Parent, time = 3 | 3.14E-01 |
| Fig. 4B | Parent, time = 3 | 1.50E-01 | Fig. S5B | Parent, time = 4 | 1.97E-01 |
| Fig. 4B | Parent, time = 4 | 1.17E-01 | Fig. S5B | Parent, time = 5 | 1.14E-01 |
| Fig. 4B | Parent, time = 5 | 1.35E-01 | Fig. S5B | Parent, time = 6 | 9.03E-02 |
| Fig. 4B | Parent, time = 6 | 1.11E-01 | Fig. S5B | Parent, time = 7 | 5.99E-02 |
| Fig. 4C | Parent, aerobic | 1.08E+01 | Fig. S5B | Parent, time = 8 | 7.13E-02 |
| Fig. 4C | Parent, microaerobic | 4.11E+00 | Fig. S5B | Parent, time = 9 | 6.07E-02 |
| Fig. 4C | MshE*, aerobic | 9.21E+00 | Fig. S5B | Parent, time = 10 | 5.96E-02 |
| Fig. 4C | MshE*, microaerobic | 3.81E+00 | Fig. S5B | Parent, time = 11 | 5.34E-02 |
| Fig. 4C | $\Delta cdpA$, aerobic | 1.23E+01 | Fig. S5B | Parent, time = 12 | 4.68E-02 |
| Fig. 4C | $\Delta cdpA$, microaerobic | 1.03E+01 | Fig. S5C | $\Delta cdpA \Delta lacZ:: cdpA-3xFLAG$, time = 0 | 1.00E+00 |
| Fig. 4D | Total CdpA, aerobic | 2.11E-01 | Fig. S5C | $\Delta cdpA \Delta lacZ:: cdpA-3xFLAG$, time = 1 | 7.02E-01 |
| Fig. 4D | Total CdpA, microaerobic | 4.20E-01 | Fig. S5C | $\Delta cdpA \Delta lacZ:: cdpA-3xFLAG$, time = 2 | 3.21E-01 |
| Fig. 4D | Upper band, aerobic | 3.52E-02 | Fig. S5C | $\Delta cdpA \Delta lacZ:: cdpA-3xFLAG$, time = 3 | 1.54E-01 |
| Fig. 4D | Upper band, microaerobic | 1.23E-01 | Fig. S5C | $\Delta cdpA \Delta lacZ:: cdpA-3xFLAG$, time = 4 | 1.13E-01 |
| Fig. 4D | Lower band, aerobic | 1.58E-01 | Fig. S5C | $\Delta cdpA \Delta lacZ:: cdpA-3xFLAG$, time = 5 | 6.35E-02 |
| Fig. 4D | Lower band, microaerobic | 2.72E-01 | Fig. S5C | $\Delta cdpA \Delta lacZ:: cdpA-3xFLAG$, time = 6 | 3.38E-02 |
| Fig. 5B | $\Delta cdpA$ - NONOate | 6.30E-01 | Fig. S5C | Parent, time = 0 | 1.00E+00 |
| Fig. 5B | $\Delta cdpA$ + NONOate | 1.03E-01 | Fig. S5C | Parent, time = 1 | 6.58E-01 |
| Fig. 5C | $\Delta cdpA$ - NONOate | 1.00E+00 | Fig. S5C | Parent, time = 2 | 3.20E-01 |
| Fig. 5C | parent - NONOate | 8.27E-01 | Fig. S5C | Parent, time = 3 | 1.70E-01 |
| Fig. 5C | $\Delta cdpA$ + NONOate | 9.33E-01 | Fig. S5C | Parent, time = 4 | 9.93E-02 |
| Fig. 5C | parent + NONOate | 8.00E-02 | Fig. S5C | Parent, time = 5 | 5.72E-02 |
| Fig. 5E | $\Delta Nap$, time = 0 | 1.00E+00 | Fig. S5C | Parent, time = 6 | 4.86E-02 |
| Fig. 5E | $\Delta Nap$, time = 1 | 5.88E-01 | | | |

**Supplemental Table 5:** Statistical Comparisons

| Comparison | P-value |  | Figure |
| --- | --- | --- | --- |
| parent (green) vs. parent (fuchsia), t = 0 | ns | >0.9999 | Fig. 1C |
| parent (green) vs. parent (fuchsia), t = 1 | ns | 0.9721 | Fig. 1C |
| parent (green) vs. parent (fuchsia), t = 2 | ns | 0.8493 | Fig. 1C |
| parent (green) vs. parent (fuchsia), t = 3 | ns | 0.6449 | Fig. 1C |
| parent (green) vs. parent (fuchsia), t = 4 | ns | 0.8493 | Fig. 1C |
| parent (green) vs. parent (fuchsia), t = 5 | ns | 0.7254 | Fig. 1C |
| parent (green) vs. parent (fuchsia), t = 6 | ns | 0.8493 | Fig. 1C |
| *pilT-mCherry* vs. parent, t = 0 | ns | >0.9999 | Fig. 1D |
| *pilT-mCherry* vs. parent, t = 1 | ns | 0.4549 | Fig. 1D |
| *pilT-mCherry* vs. parent, t = 2 | * | 0.0380 | Fig. 1D |
| *pilT-mCherry* vs. parent, t = 3 | *** | 0.0005 | Fig. 1D |
| *pilT-mCherry* vs. parent, t = 4 | **** | <0.0001 | Fig. 1D |
| *pilT*-mCherry vs. parent, t = 5 | **** | <0.0001 | Fig. 1D |
| *pilT*-mCherry vs. parent, t = 6 | **** | <0.0001 | Fig. 1D |
| *P*<sub>BAD</sub>-dcpA Δ*vps* vs. parent, t = 0 | ns | >0.9999 | Fig. 2A |
| *P*<sub>BAD</sub>-dcpA Δ*vps* vs. parent, t = 1 | ns | 0.2476 | Fig. 2A |
| *P*<sub>BAD</sub>-dcpA Δ*vps* vs. parent, t = 2 | *** | 0.0001 | Fig. 2A |
| *P*<sub>BAD</sub>-dcpA Δ*vps* vs. parent, t = 3 | **** | <0.0001 | Fig. 2A |
| *P*<sub>BAD</sub>-dcpA Δ*vps* vs. parent, t = 4 | **** | <0.0001 | Fig. 2A |
| *P*<sub>BAD</sub>-dcpA Δ*vps* vs. parent, t = 5 | **** | <0.0001 | Fig. 2A |
| *P*<sub>BAD</sub>-dcpA Δ*vps* vs. parent, t = 6 | **** | <0.0001 | Fig. 2A |
| *mshE* * vs. parent, t = 0 | ns | >0.9999 | Fig. 2B |
| *mshE* * vs. parent, t = 1 | ns | 0.4192 | Fig. 2B |
| *mshE* * vs. parent, t = 2 | ** | 0.0062 | Fig. 2B |
| *mshE* * vs. parent, t = 3 | **** | <0.0001 | Fig. 2B |
| *mshE* * vs. parent, t = 4 | **** | <0.0001 | Fig. 2B |
| *mshE* * vs. parent, t = 5 | **** | <0.0001 | Fig. 2B |
| *mshE* * vs. parent, t = 6 | **** | <0.0001 | Fig. 2B |
| Δ*cdpA*, Δ*lacZ*::*cdpA* vs. parent, time = 0 | ns | >0.9999 | Fig. 4A |
| Δ*cdpA*, Δ*lacZ*::*cdpA* vs. parent, time = 1 | ns | 0.9778 | Fig. 4A |
| Δ*cdpA*, Δ*lacZ*::*cdpA* vs. parent, time = 2 | ns | 0.9309 | Fig. 4A |
| Δ*cdpA*, Δ*lacZ*::*cdpA* vs. parent, time = 3 | ns | 0.8906 | Fig. 4A |
| Δ*cdpA*, Δ*lacZ*::*cdpA* vs. parent, time = 4 | ns | 0.8244 | Fig. 4A |
| Δ*cdpA*, Δ*lacZ*::*cdpA* vs. parent, time = 5 | ns | 0.8864 | Fig. 4A |
| Δ*cdpA*, Δ*lacZ*::*cdpA* vs. parent, time = 6 | ns | 0.3401 | Fig. 4A |
| Δ*cdpA*, Δ*lacZ*::*cdpA*<sup>ECL→AAA</sup> vs. parent, time = 0 | ns | >0.9999 | Fig. 4B |
| Δ*cdpA*, Δ*lacZ*::*cdpA*<sup>ECL→AAA</sup> vs. parent, time = 1 | ns | 0.1993 | Fig. 4B |
| Δ*cdpA*, Δ*lacZ*::*cdpA*<sup>ECL→AAA</sup> vs. parent, time = 2 | ** | 0.0032 | Fig. 4B |
| Δ*cdpA*, Δ*lacZ*::*cdpA*<sup>ECL→AAA</sup> vs. parent, time = 3 | **** | <0.0001 | Fig. 4B |
| Δ*cdpA*, Δ*lacZ*::*cdpA*<sup>ECL→AAA</sup> vs. parent, time = 4 | **** | <0.0001 | Fig. 4B |
| Δ*cdpA*, Δ*lacZ*::*cdpA*<sup>ECL→AAA</sup> vs. parent, time = 5 | **** | <0.0001 | Fig. 4B |
| Δ*cdpA*, Δ*lacZ*::*cdpA*<sup>ECL→AAA</sup> vs. parent, time = 6 | **** | <0.0001 | Fig. 4B |
| parent, aerobic vs. microaerobic | ** | 0.0025 | Fig. 4C |
| *mshE* *, aerobic vs. microaerobic | ** | 0.0088 | Fig. 4C |
| Δ*cdpA*, aerobic vs. microaerobic | ns | 0.2484 | Fig. 4C |
| Total CdpA, aerobic vs. microaerobic | ns | 0.0773 | Fig. 4D |
| Upper band, aerobic vs. microaerobic | ** | 0.0015 | Fig. 4D |
| Lower band, aerobic vs. microaerobic | ns | 0.1172 | Fig. 4D |
| Δ*cdpA* vs. Δ*cdpA* + NONOate | **** | <0.0001 | Fig. 5B |
| parent vs. Δ*cdpA*, - NONOate | ns | 0.8180 | Fig. 5C |
| parent vs. Δ*cdpA*, + NONOate | **** | <0.0001 | Fig. 5C |
| Δ*Nap* vs. parent, time = 0 | ns | >0.9999 | Fig. 5E |
| Δ*Nap* vs. parent, time = 1 | ns | 0.9221 | Fig. 5E |
| Δ*Nap* vs. parent, time = 2 | ns | 0.8995 | Fig. 5E |
| Δ*Nap* vs. parent, time = 3 | ns | 0.8995 | Fig. 5E |
| Δ*Nap* vs. parent, time = 4 | ns | 0.9973 | Fig. 5E |
| Δ*Nap* vs. parent, time = 5 | ns | 0.9539 | Fig. 5E |
| Δ*Nap* vs. parent, time = 6 | ns | 0.9221 | Fig. 5E |
| Δ*vps* vs. parent, time = 0 | ns | >0.9999 | Fig. S1C |
| Δ*vps* vs. parent, time = 1 | ns | 0.9873 | Fig. S1C |
| Δ*vps* vs. parent, time = 2 | ns | 0.8283 | Fig. S1C |
| $\Delta vps$ vs. parent, time = 3 | ns | 0.9873 | Fig. S1C |
| $\Delta vps$ vs. parent, time = 4 | ns | 0.9235 | Fig. S1C |
| $\Delta vps$ vs. parent, time = 5 | ns | 0.8283 | Fig. S1C |
| $\Delta vps$ vs. parent, time = 6 | ns | 0.3354 | Fig. S1C |
| $P_{BAD-cdgJ}$ vs. $P_{BAD-cdgJ} mshE^*$ | **** | <0.0001 | Fig. S3A |
| $P_{BAD-cdgJ}$ vs. $\Delta MSHA$ | **** | <0.0001 | Fig. S3A |
| parent, aerobic vs. microaerobic | *** | 0.0004 | Fig. S4A |
| $mshE^*$, aerobic vs. microaerobic | ns | 0.1874 | Fig. S4A |
| $\Delta cdpA$, aerobic vs. microaerobic | ns | 0.9463 | Fig. S4A |
| $\Delta cdpA$ vs. parent, time = 0 | ns | >0.9999 | Fig. S5B |
| $\Delta cdpA$ vs. parent, time = 1 | ns | 0.9139 | Fig. S5B |
| $\Delta cdpA$ vs. parent, time = 2 | ns | 0.1646 | Fig. S5B |
| $\Delta cdpA$ vs. parent, time = 3 | **** | <0.0001 | Fig. S5B |
| $\Delta cdpA$ vs. parent, time = 4 | **** | <0.0001 | Fig. S5B |
| $\Delta cdpA$ vs. parent, time = 5 | **** | <0.0001 | Fig. S5B |
| $\Delta cdpA$ vs. parent, time = 6 | **** | <0.0001 | Fig. S5B |
| $\Delta cdpA$ vs. parent, time = 7 | **** | <0.0001 | Fig. S5B |
| $\Delta cdpA$ vs. parent, time = 8 | **** | <0.0001 | Fig. S5B |
| $\Delta cdpA$ vs. parent, time = 9 | **** | <0.0001 | Fig. S5B |
| $\Delta cdpA$ vs. parent, time = 10 | **** | <0.0001 | Fig. S5B |
| $\Delta cdpA$ vs. parent, time = 11 | **** | <0.0001 | Fig. S5B |
| $\Delta cdpA$ vs. parent, time = 12 | **** | <0.0001 | Fig. S5B |
| $\Delta cdpA$, $\Delta lacZ::cdpA-3xFLAG$ vs. parent, time = 0 | ns | >0.9999 | Fig. S5C |
| $\Delta cdpA$, $\Delta lacZ::cdpA-3xFLAG$ vs. parent, time = 1 | ns | 0.9953 | Fig. S5C |
| $\Delta cdpA$, $\Delta lacZ::cdpA-3xFLAG$ vs. parent, time = 2 | ns | 0.9990 | Fig. S5C |
| $\Delta cdpA$, $\Delta lacZ::cdpA-3xFLAG$ vs. parent, time = 3 | ns | 0.9976 | Fig. S5C |
| $\Delta cdpA$, $\Delta lacZ::cdpA-3xFLAG$ vs. parent, time = 4 | ns | 0.9151 | Fig. S5C |
| $\Delta cdpA$, $\Delta lacZ::cdpA-3xFLAG$ vs. parent, time = 5 | ns | 0.9725 | Fig. S5C |
| $\Delta cdpA$, $\Delta lacZ::cdpA-3xFLAG$ vs. parent, time = 6 | ns | 0.3426 | Fig. S5C |

